# Two copy-number forms of the *Kala4/OsB2* upstream duplication in purple rice revealed by a telomere-to-telomere genome of landrace ZN65

**DOI:** 10.64898/2026.07.07.737121

**Authors:** Feng Zhao, Juan Zhao, Fang Zhao, Suhong Bai, Yahan Wu, Rengguo Zhu

**Affiliations:** Key Laboratory of Ethnic Medical Resources Research and Southeast Asian International Cooperation in Yunnan Province, School of Tea and Coffee and School of Bioinformatics and Engineering, Pu’er University, 6 Xueyuan Road, Pu’er, Yunnan 665000, China; Yunnan International Joint Laboratory of Digital Conservation and Germplasm Innovation and Application of China-Laos Tea Resources, Pu’er University, Pu’er, Yunnan 665000, China

**Keywords:** Telomere-to-telomere genome, Pigmented rice, Anthocyanin, *Kala4*/*OsB2*, *OsKala3*, Structural variation, Copy-number variation

## Abstract

**Background:** Black and purple pericarp in rice requires activation of the regulator *Kala4*/*OsB2*. The canonical black-rice allele is a promoter rearrangement that was described from DNA gel blots and amplified fragments, and its sequence is unavailable. Whether pigmented landraces carry one upstream structure or several is therefore unknown. We examined this region in the purple-pericarp landrace Mojiang ZN65 and its distribution in sequenced rice germplasm.

**Results:** We assembled a gap-free telomere-to-telomere genome of ZN65 (395.1 Mb; all 24 telomeres). Upstream of *Kala4*/*OsB2*, a 5.87-kb segment (block A) and a 6.0-kb segment that contains the first two annotated exons of *OsB2* were each present in three non-tandem copies. The arrangement was reproduced by an independent HiFi-only assembly and was spanned by three Nanopore reads. Among 2,913 accessions of the 3,000 Rice Genomes panel, read depth detected block A gain in 29 (1.0%), 17 at three or more copies and 12 at two. All 29 carried read pairs joining block A to the *OsB2* 5′ segment, and none of 58 matched controls did. A junction specific to the third copy was found in 16 of these 17 and in 2 of these 12. Among 2,055 accessions with recorded seed-coat colour, 18 of 19 purple accessions carried the gain, and purple was recorded for 9 of 13 and 9 of 11 carriers of the two classes. Among 210 NCBI assemblies and 264 Genome Warehouse materials, five carried the junctions with two copies of block A, and one, the Chinese accession Momi, carried three copies, which its Nanopore reads confirmed. An elevated *OsKala3* promoter-repeat depth ratio was found in 21 of 29 carriers, and four purple carriers lacked it.

**Conclusions:** ZN65 provides a gap-free, read-supported sequence of a *Kala4*/*OsB2* upstream duplication that occurs in at least two copy-number forms in purple rice. Its gene-proximal part has the same kinds of elements in the same order as the published description of the canonical black-rice allele, although the segment lengths differ. Both forms were found in purple accessions, and the additional copy showed no relation to recorded colour. Its effect on *OsB2* expression remains untested.

## Background

Black, purple and red rice landraces are valued for the anthocyanins and proanthocyanidins that accumulate in the grain pericarp and are a reservoir of nutritional and agronomic diversity [1]. Pericarp anthocyanin pigmentation is controlled chiefly by three loci: *Kala1*, the biosynthetic gene *OsDFR*; *Kala3*, the R2R3-MYB *OsKala3*; and *Kala4*, the basic helix-loop-helix (bHLH) protein *OsB2*. The two regulators form the MYB-bHLH-WD40 (MBW) complex with the WD40 protein *OsTTG1* and activate the anthocyanin structural genes (gene nomenclature in Additional file 2: Table S8) [2, 3]. This pericarp system is genetically distinct from the *C*–*S*–*A* system that colours the hull and apiculus [4].

The black pericarp is a gain of function caused by rearrangement of the *Kala4*/*OsB2* promoter, which drives ectopic *OsB2* expression in the pericarp [2]. The canonical black-rice allele combines a duplicated segment with a large upstream insertion, and its variants define distinct promoter types among black cultivars [2]. It was characterized by DNA gel blots and polymerase chain reaction, and that study lists no sequence accession for the rearranged region. A second regulatory variant lies at *Kala3*: black-pericarp cultivars carry two copies of a promoter repeat unit where white- and red-pericarp cultivars carry one, and repeat copy number scales with transactivation strength [3]. Both variants lie in repeat-rich sequence that is difficult to resolve in fragmented assemblies. Gap-free assemblies now make such loci tractable [5, 6]. A gain-of-function *Kala4* haplotype was recently reported in black accessions of a Vietnamese landrace panel [7], but the sequence of the rearranged region remains unreported. It is therefore unknown whether pigmented landraces carry one upstream structure or several.

We addressed this question with Mojiang ZN65, a Hani purple-pericarp landrace from Yunnan, China. The central question was what structure ZN65 carries upstream of *Kala4*/*OsB2*, and whether purple rice carries this structure in one form or in several. It was divided into three questions:

1. Q1 (structure): the structure of the *Kala4*/*OsB2* upstream region in ZN65, whether it is supported by evidence independent of the assembly, and whether public assemblies carry it.
2. Q2 (distribution): the frequency and copy-number forms of gain of the ZN65 upstream block (block A) in the 3,000 Rice Genomes panel, whether the accessions with that gain carry the ZN65 junctions and a shared flanking genotype, and whether the gain tracks recorded seed-coat colour.
3. Q3 (relation to *OsKala3*): the distribution of the gain relative to the duplicated *OsKala3* promoter repeat.

To answer them, we assembled a gap-free telomere-to-telomere (T2T) genome of ZN65, examined the *Kala4*/*OsB2* region with additional read-based checks, and surveyed 3,011 accessions of the 3,000 Rice Genomes panel by read depth, junction-spanning reads and flanking genotypes. We also screened the 210 *Oryza sativa* assemblies listed at NCBI and the 466 rice assemblies in the Genome Warehouse for block A and the junctions of the structure.

## Methods

### Plant material and sequencing

All genome libraries were prepared from a single plant of ZN65 (provenance and voucher under Declarations). Genomic DNA was sequenced on PacBio Revio (high-fidelity [HiFi] circular consensus sequencing, CCS; 20.6 Gb, read N50 19.4 kb), Oxford Nanopore Technologies (ONT; ultra-long; 256,971 reads, 21.68 Gb, read N50 100 kb; base-called with Guppy, mean Q-score ≥7), DNBSEQ (72.6 Gb clean) and Hi-C (39.3 Gb clean). Reads were quality-filtered with fastp v0.19.5.

### Genome assembly and annotation

Contigs were assembled from the ONT ultra-long reads alone with hifiasm v0.25.0 (ONT mode) [8]. HiFi reads were excluded from contig construction. Contigs were error-corrected with NextPolish v1.4.1 using HiFi and DNBSEQ reads and scaffolded to twelve chromosomes with Juicer v1.6 and 3D-DNA using Hi-C. Gaps were filled and telomeres identified with quarTeT v1.2.5 [9]. Chromosome 4 contained no gap before this step. The assembly was polished by mapping HiFi/ONT with Winnowmap v2.03 and DNBSEQ with bwa v0.7.12, followed by Pilon v1.24 (`--fix snps,indels -- minmq 10`). The final assembly is a single collapsed representation of twelve chromosomes. Quality was assessed with Merqury v1.3 (k = 19) [10], the long terminal repeat (LTR) Assembly Index (LAI; LTR_retriever) [11], read-mapping rates and Benchmarking Universal Single-Copy Orthologues (BUSCO) v5.8.0 (embryophyta_odb10) [12]. Centromeres were localized as *CentO* satellite arrays (Tandem Repeats Finder monomers of 155–165 bp) and the 45S/5S rDNA arrays from the rRNA annotation (Additional file 2: Table S5). Repeats were annotated with RepeatModeler v2.0.6 [13], LTR_FINDER v1.07, RepeatMasker/RepeatProteinMask (RepBase) and Tandem Repeats Finder v4.09.1. Protein-coding genes were predicted by integrating three lines of evidence with EVidenceModeler v2.1.0: *ab initio* prediction (AUGUSTUS v3.2.3), protein homology (Miniprot v0.15, proteomes of eight species) and transcript alignments (PASA v2.4.1, rice cDNA and expressed sequence tag collections). Non-coding RNAs were annotated with tRNAscan-SE v2.0.7, BLASTn (rRNA) and Rfam/cmsearch (miRNA, snRNA). Gene function was assigned with Diamond v2.1.10 (E ≤ 10^−5^) against NR, SwissProt, TrEMBL and the Kyoto Encyclopedia of Genes and Genomes (KEGG), and with InterProScan.

### Comparative genomics and structural variation

ZN65 was aligned to Nipponbare (IRGSP-1.0), to the gap-free *indica* references MH63RS3 and ZS97RS3 [5] and to the assembly registered as Cempo-Ireng (GenBank GCA_055776245.1) with minimap2 (asm5) [14]. Structural variants were called with SyRI [15] and visualized with plotsr [16]. Counts refer to SyRI per-event-type annotations, in which translocations and inverted translocations, and duplications and inverted duplications, are tabulated separately. Robustness was checked by repeating the ZN65–Nipponbare comparison at the asm10 and asm20 presets (121, 132 and 126 inversions at asm5/asm10/asm20; the major inversions, including the chromosome-6 inversion, were recovered at all three) and by nucmer/Assemblytics [17]. Nipponbare-nonaligned sequence (73.2 Mb) was the sum of the SyRI not-aligned (NOTAL; 70.3 Mb) and insertion (INS; 2.9 Mb) categories (Additional file 2: Table S3). Because it is defined against a *japonica* reference, it conflates subspecies divergence with ZN65-specific gain. To separate the two, the NOTAL component was intersected with alignments of ZN65 to MH63RS3 and ZS97RS3 (minimap2 -c -x asm10, --secondary=no). INS was excluded from this intersection because SyRI reports insertions as breakpoints and gives no query intervals for them. Neither block A nor the *OsKala3* promoter repeat unit (RU) falls within NOTAL, as expected for copy-number gains of sequence that is single-copy in Nipponbare. Pigmentation loci were extracted with their flanks and compared with nucmer/dnadiff [18] and MAFFT [19]. Orthologues were identified by reciprocal BLAST against the IRGSP-1.0 proteome, and orthogroups with OrthoFinder v2.5 (Diamond) [20].

### Initial block A screen across 34 assemblies

Qualifying homologous hits to the ∼5.87-kb block A were counted across 34 distinct assemblies (36 files screened; MH63RS3 and GCA_016134885.1, and ZS97RS3 and GCA_001623345.3, were identical across all twelve nuclear chromosomes and each pair was counted once), spanning cultivated *Oryza sativa* (*indica*, *japonica*, *aus*, tropical *japonica*), *O. rufipogon*, *O. nivara*, *O. longistaminata*, *O. meridionalis*, *O. australiensis*, *O. glaberrima* and *O. barthii* (Additional file 2: Table S10). Nipponbare and N22 are each represented by two distinct assemblies. Hits were called by whole-genome alignment (minimap2 -c -x asm20 -N 20 -p 0), retaining alignment blocks ≥1 kb and merging blocks on the same target sequence and strand into collinear clusters when the target gap was non-negative and <5 kb, the absolute query gap was <5 kb and the query overlap was <25% of the shorter block’s query span. Each cluster covering ≥50% of the 5,870-bp query was counted as one hit. Cluster identity was the sum of matching bases divided by the summed alignment-block length, and query coverage the union of query intervals divided by the query length. The search was not calibrated for detection sensitivity, so counts are lower bounds on copy number. Hits below 95% identity or 95% query coverage are reported as coverage- or divergence-limited.

### Validation of the *Kala4*/*OsB2* region

The two composite gains were delimited in ZN65 coordinates from SyRI insertions relative to Nipponbare and corroborated by nucmer --maxmatch. HiFi reads were mapped with minimap2 (map-hifi). A read was counted as spanning an interval if it was a primary alignment with mapping quality (MAPQ) ≥ 50, carried no supplementary-alignment (SA) tag, extended ≥1 kb beyond both ends of the interval and contained no ≥1-kb indel within it. Oxford Nanopore ultra-long reads spanning the whole three-copy region were identified in the same way (reads 04d6d3c8, f4924c3f and dade46d3; MAPQ 60). Four further checks were made. First, the HiFi reads, which were held out of contig construction, were assembled alone with hifiasm (default HiFi mode), and the 250-kb region around the locus was aligned to the resulting contigs with minimap2 (asm5). Second, structural assembly errors were called genome-wide with Inspector [21] from the HiFi alignments. Third, the ZN65 DNBSEQ reads (first 40 million pairs, ∼32×) were mapped to Nipponbare with bwa-mem2 and processed with the read-depth and junction procedures used for the population survey, as a positive control for those procedures. Fourth, Hi-C read pairs were mapped with chromap (MAPQ ≥ 10) and contacts between the copies were compared with distance-matched background intervals. Heterozygosity was assessed from the 21-mer spectrum of the DNBSEQ reads (KMC and GenomeScope 2.0) [22] and from heterozygous single-nucleotide polymorphisms (SNPs) called from the HiFi alignments with bcftools, genome-wide and within chr4:27.99–28.16 Mb. The three block A copies and their 2-kb flanks were compared pairwise with minimap2.

### Phylogenetics, pathway inventory and dating

Protein alignments were built with MAFFT (L-INS-i) [19] and maximum-likelihood trees inferred with IQ-TREE 2 (ModelFinder; 1,000 ultrafast bootstraps) [23]. Pathway genes were assigned to enzymatic steps by KEGG orthology in ZN65, and the set was extended to all ZN65 genes and Nipponbare loci in the same orthogroups. Each orthogroup was assigned to the step with the most KEGG-assigned ZN65 genes, so that each ZN65 gene and each Nipponbare locus was counted once. To test for missed annotations, the protein of every Nipponbare pathway locus was aligned to the ZN65 genome with miniprot [24], and each best alignment was classified by whether it overlapped an annotated ZN65 gene and whether it was intact (query coverage of at least 0.9 with no frameshift or premature stop codon). The insertion age of the intronic *Gypsy* element was estimated from the Kimura two-parameter (K2P) distance *K* between its two LTRs (aligned with EMBOSS stretcher; gap sites excluded) as *T* = *K*/(2µ), with µ = 1.3 × 10^−8^ substitutions per site per year [25]. A 95% confidence interval (CI) was obtained from 1,000 bootstrap replicates of alignment columns.

### Subspecies placement

ZN65 was genotyped at the 3,000 Rice Genomes Project pruned SNP set (1,011,601 SNPs) [26] from its whole-genome alignment to Nipponbare, merged into the 3,024-accession panel with PLINK 1.9 and projected by principal-component analysis (PLINK 2.0). Subpopulation labels were taken from the published K = 9 admixture matrix, and accessions with a maximum ancestry coefficient below 0.65 were labelled admixed.

### Population copy-number survey at block A

Public alignments to Nipponbare (IRGSP-1.0) of the 3,000 Rice Genomes panel [26] were queried by remote range requests in three batches (558, 1,264 and 1,189 accessions; 3,011 in total). For each accession, reads were counted with samtools view (`-c -q 20`) [27] over a counting window on block A at its orthologous position (chr04:28,022,288–28,028,159; 5,872 bp, starting one base upstream of the block A boundary) and two size-matched single-copy flanking controls (chr04:27,900,000–27,905,871 and chr04:28,200,000–28,205,871). The depth ratio (block A read density divided by the mean density of the two controls) was calibrated within each batch by setting the batch median ratio (1.188, 1.137 and 1.113) to one copy. Accessions were called from this calibrated estimate as one copy (estimate <1.45), two copies (1.45 ≤ estimate <2.45) or three or more (≥2.45). Accessions were excluded when an interval returned no count, when a control interval returned fewer than 25% of the batch median control count, or when the two control counts differed by more than threefold, leaving 2,913 (Additional file 2: Table S12). For all accessions called at two or more copies and for two single-copy controls per carrier, matched by subpopulation and control read count, reads over chr04:28,012,000–28,038,000 were retrieved. Depth in 500-bp bins was divided by the accession’s control density and by the median of the 58 controls in the same bin. Bins in which the control median fell below 0.7 of the control density were discarded. Reads over chr04:27,925,000–27,950,000, which contains the 5′ end of *OsB2*, were retrieved for the same accessions and processed in the same way. The relative depth of each accession was averaged over the retained bins within the 6.0-kb segment of I1 (bin starts 27,935,000–27,940,000). For two accessions with elevated depth beyond block A, starts of alignments in chr04:27.80–28.30 Mb were counted in 5-kb bins, divided by the accession’s median outside 27.90–28.05 Mb and by the median of eight matched controls in the same bin. Bins with a control density below 0.5 were discarded.

### Junction analysis

The junction positions were first defined in ZN65 itself. ZN65 DNBSEQ reads mapped to Nipponbare were scanned over chr04:28,012,000–28,038,000 for soft-clipped reads (clip ≥20 bp, primary alignments). Clip positions were clustered within 30 bp and clusters with ≥3 reads retained. The clipped segments of each cluster were aligned to Nipponbare and to the ZN65 assembly (minimap2 `-c -k11 -w5 -x sr`), and the whole reads to the ZN65 assembly, to separate junctions of the locus from reads of paralogous origin. Three signatures of the ZN65 structure were obtained in Nipponbare coordinates: a left clip at 28,022,289 whose clipped segment ends at 27,940,807 (J1), a right clip at 28,030,604 whose clipped segment starts at 27,934,782 (J2), and a right clip at 28,029,146 whose clipped segment is repetitive (J3; specific to the second copy, A2). In the panel, reads clipped at these three positions (±5 bp) were counted for each carrier and control, and clipped segments were aligned to Nipponbare genome-wide to test the expected partner position (±2 bp). Discordant pairs were counted as read pairs with one end inside block A and the mate at 27.930–27.945 Mb.

### Public assemblies

All 210 assemblies of *Oryza sativa* listed by the NCBI Datasets service on 21 November 2026 were screened. Sequences were retrieved by accession from the European Nucleotide Archive or from NCBI. Each file was accepted when its total length equalled the length registered at NCBI or when its checksum matched the NCBI record. Block A and five 600-bp probes were aligned to each assembly with minimap2 (asm5). Four probes were centred on ZN65 junctions (the two junctions of I1, the corresponding junction of the third copy, and an A2-specific position), and one was centred on a reference-like position at the left end of I2 and served as a control for coverage of the region. A block A hit required ≥95% query coverage and ≥98% identity. A probe was scored as present when one alignment extended at least 150 bp on both sides of its centre at ≥97% identity with no indel of 5 bp or more overlapping the central 300 bp. An assembly in which the control probe was absent was treated as not evaluable. The 135-kb ZN65 locus (Chr4:28,010,000–28,145,000) was aligned to each junction-positive assembly in the same way. The 6.0-kb segment of I1 (the Nipponbare sequence chr04:27,934,782–27,940,807 and the ZN65 copy contiguous with the gene), the 8.3-kb segment that follows it in ZN65, block A, I2 and the rest of *OsB2* were aligned to ZN65, to the junction-positive assemblies and to Copsemar9 with minimap2 (asm5). Alignments that covered at least 90% of a segment were recorded (Additional file 2: Table S15). Exon positions of *OsB2* in Nipponbare were taken from transcript Os04t0557500-01 of the IRGSP-1.0 annotation (Ensembl Plants). Pericarp colour was taken from Sedeek et al. [28] for the five varieties of that study, because the sample records of the assemblies omit it.

The two Cempo assemblies were compared with the HiFi reads of Sedeek et al. [28] (BioProject PRJNA942452; SRR23822209, deposited as Cempo Ireng; SRR23822210, deposited as Cempo Abang), which were retrieved from the European Nucleotide Archive and verified by checksum. The first 20,000 reads of each run were aligned to both assemblies with minimap2 (map-hifi). The first 20,000 reads of the other three varieties (SRR23822207, Pulut Hitam; SRR23822211, Zag; SRR23822212, Balatinaw) were read from the start of each run file, and the reads of all five runs were aligned to the five assemblies in the same way. For primary alignments of at least 5 kb with a mapping quality of at least 20, divergence was calculated as the summed edit distance divided by the summed alignment length. All reads of the two Cempo runs were also aligned to the five probes and to Nipponbare. We counted the reads that covered positions 31–570 of a probe in one alignment with less than 1% divergence, and compared the mean depth of block A with the median depth of the 100-kb intervals on both sides. At the site of the 14-bp deletion that defines the *rc* allele of Nipponbare [29], we counted the spanning reads that carried a 14-bp insertion with the sequence of the functional allele or a rotation of it, as placed by the aligner. The same site was examined in the assemblies.

Assemblies of *Oryza* in the Genome Warehouse (GWH) [30] were screened in the same way. The genome files under the *Oryza* entries of the GWH download server were listed on 23 November 2026 (3,221 files). The 2,755 files of two BioProjects (PRJCA017583 and PRJCA017591), which deposit microbial isolate and metagenome-assembled genomes under these entries, were excluded, which left 466 rice assemblies. Each file was checked against its GWH md5 checksum (262 assemblies) or, when no checksum was listed, by its registered size and the gzip integrity test (204 assemblies). Versions of one accession and assemblies of one variety were grouped as one material according to the GWH sample records and the accession table of Shang et al. [31], which gave 264 materials. The locus and segment alignments were repeated for the junction-positive assemblies.

The Nanopore reads of NH202 (SRR12451611; BioProject PRJNA656318; 1,426,900 reads) [31] were retrieved from NCBI, verified by checksum and aligned with minimap2 (map-ont) to ZN65 Chr4:27,950,001–28,250,000. For each gain, two anchor positions outside it were used (Chr4:28,016,000 and 28,037,000 for I1; Chr4:28,117,800 and 28,140,200 for I2). For reads with alignments of mapping quality ≥20 that covered both anchors, the distance between the anchors on the read was compared with that in ZN65 (21,000 bp for I1 and 22,400 bp for I2) and, for I2, with the distance expected without I2 (8,643 bp). A read was scored as three-copy-like when the distance was within 10% of the ZN65 value and as two-copy-like when it was within 15% of 8,643 bp.

### Seed-coat colour

Seed-coat colour was taken from the morpho-agronomic records of the 3,000 Rice Genomes panel (descriptor SCCO_REV), which were scored at the International Rice Research Institute. Codes 010, 050–055, 070 and 080 were read as white, brown, red and purple. One record with code 088 was read as purple, and five records with code 999 were kept as a class other than purple; the phenotype file of the panel defines code 088 as variable purple and, in its code list for records before 2007, code 999 as mixture, and the association was also recomputed with these six records treated as missing.

### Flanking genotypes

Genotypes at the pruned SNP set were extracted for chr04:26–30 Mb, and SNPs within the duplicated segments (chr04:27,934,782–27,940,807 and 28,022,289–28,030,604) were removed. Pairwise genotype distance was the mean absolute difference in allele dosage divided by two, over the SNPs within 200 kb of the locus. The shared segment of a pair was measured outward from the left boundary of block A to the first SNP at which the two accessions were homozygous for different alleles on each side. Missing and heterozygous genotypes were not counted as conflicts. Observed means were compared with 2,000 random sets of non-carriers drawn with the same subpopulation composition. The data are unphased, so these measures describe genotype similarity and do not establish identical haplotypes.

### *OsKala3* identification and promoter repeats

The ZN65 *OsKala3* orthologue was identified by DIAMOND blastp against the ZN65 protein set and three criteria: location on chromosome 3, the subgroup 6 (SG6)-like motif xPxPRTx reported to be absent from *OsC1*, and the subgroup 5 (SG5) DNEI motif [3]. The 6 kb upstream of the coding sequence (CDS) start (minus strand; start codon ATG at chr03:17,954,449) was reverse-complemented and scanned for tandem repeats by blastn self-alignment and by an independent periodicity scan (period 300–360 bp, identity >85%). The Nipponbare orthologue, located by splice-aware alignment of the ZN65 CDS (966/966 matched), was processed identically.

### RU dosage

RU read-depth ratios were obtained as for block A, using the RU interval chr03:16,883,626–16,883,960 (located by aligning the ZN65 RU to Nipponbare at 100% identity) and two equally sized single-copy controls (chr03:16,760,000–16,760,334 and chr03:17,010,000–17,010,334). The ratio was the RU count divided by the mean of the two control counts, and accessions with a mean control count below 20 reads were excluded. A ratio of 1.5 or more was taken as elevated, and thresholds from 1.3 to 2.0 were examined.

### Statistics

Association between block A gain (two or more copies) and purple seed coat was examined in the 2,055 accessions with a copy-number call, a colour record and genotypes. We report the contingency tables, two-sided Fisher’s exact tests, an exact conditional test stratified by subpopulation (Mantel–Haenszel, R 4.6.1), Firth-penalized logistic regression [32] with the first ten principal components as covariates (PLINK 2.0), and a logistic mixed model with a genomic relationship matrix as random effect (GMMAT v1.5.0; score test, without and with the first four principal components) [33]. In the Firth regression, carrier status was coded 0 or 2, and the odds ratio for carriers is the square of the estimate per unit (36.8). A genome-wide scan was also run with a linear mixed model (GEMMA v0.98.5; centred relatedness matrix; Wald test) over 746,091 SNPs, with carrier status added as one marker. The association was also examined within accessions whose flanking genotype lay within a fixed distance of the modal genotype of the three-copy carriers. Carriers share a flanking genotype and are not independent observations, so all *P* values are reported as exploratory.

### Use of large language models

Claude (Anthropic) was used to help write and debug analysis and plotting scripts and to edit the English text. All analyses were run on the authors’ computers, and all results, numbers and conclusions were checked by the authors against the underlying data files. The authors take full responsibility for the content.

## Results

### A gap-free telomere-to-telomere assembly of ZN65

The ZN65 assembly comprises twelve gap-free chromosomes totalling 395,112,834 bp (contig N50 32.35 Mb; GC 43.66%), and all 24 chromosome ends carry the plant telomere repeat (AAACCCT) (Figure 1). Consensus quality value was 53.6 (base error rate 4.4 × 10^−6^), LAI 15.5, BUSCO completeness 99.6% (1,607 of 1,614) and read-mapping rates 99.31–99.99% (Additional file 2: Table S1). Inspector reported a single structural error genome-wide, a 55-bp collapse on chromosome 5. HiFi alignments supported 528 heterozygous SNPs genome-wide, whereas the 21-mer spectrum gave a heterozygosity estimate of 0.21–0.23%. This difference remains unresolved, and the sequenced plant is treated as largely homozygous on the basis of the direct count. All twelve centromeres were localized as *CentO* arrays, and the 45S rDNA arrays on chromosomes 9 and 10 and the 5S rDNA array on chromosome 11 were assembled (Additional file 2: Table S5). A Fourier scan of the assembly located a satellite array with a period of 140–170 bp that overlapped the centromere core on each of the twelve chromosomes (Additional file 1: Figure S3). The median peak period of these arrays was 154.8–155.7 bp on chromosomes 2, 3, 4, 5 and 7, 153.8 bp on chromosome 8, 164.3 and 164.9 bp on chromosomes 9 and 10, and 158.9 bp on chromosome 1. The windows of chromosome 11 peaked at 154.4–162.6 bp. On chromosomes 6 and 12, two windows peaked at 140.6–140.7 and 149.6 bp, respectively, and the others at 150.2–153.2 and 155.5–161.3 bp. The scan also detected arrays of a 353-bp satellite at the long-arm ends of chromosomes 5, 10, 11 and 12, whose distal boundaries Tandem Repeats Finder placed 15–27 kb from the chromosome ends. Annotation identified 42,090 protein-coding genes (Additional file 2: Table S2). Repeats occupied 56.6% of the genome and were concentrated in pericentromeric regions.

**Figure 1.**
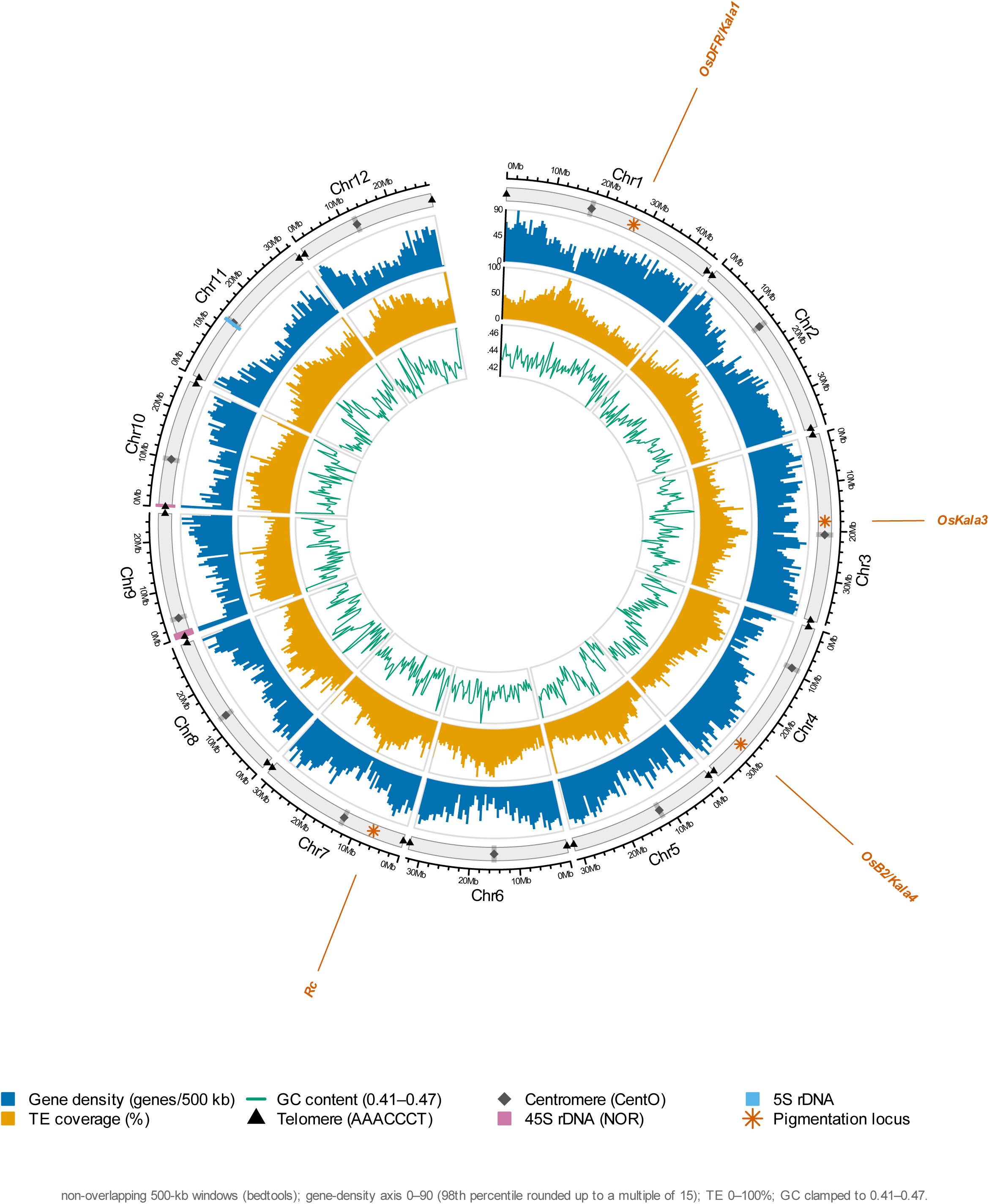
Telomere-to-telomere genome landscape of the purple rice landrace ZN65. Rings, outer to inner: chromosome ideogram with 10-Mb ticks (filled triangles, resolved telomeres [24/24]; diamonds, *CentO*-satellite centromeres; coloured bars, 45S [pink] and 5S [blue] rDNA arrays; asterisks with leader lines, the four pericarp-pigmentation loci *OsB2*/*Kala4*, *OsDFR*/*Kala1*, *OsKala3* and *Rc*); gene density (genes per 500 kb; axis 0–90, the 98th percentile rounded up to a multiple of 15); transposable-element (TE) coverage (0–100%); GC content (line; axis 0.41–0.47). All tracks use non-overlapping 500-kb windows (bedtools).

Relative to the *japonica* reference Nipponbare, ZN65 carried 1,045,956 single-nucleotide variants and 121 inversions, including a large inversion on chromosome 6, among other structural variants (Figure 2; Additional file 2: Table S3), and was larger on every chromosome (∼22 Mb in total). Of the 73.2 Mb of ZN65 sequence without an aligned Nipponbare counterpart, 45.1 Mb (62%) was repetitive, mostly LTR/*Gypsy* (26.9 Mb). However, 91.4% of the 70.3-Mb not-aligned (NOTAL) component was present in at least one of the gap-free *indica* references MH63RS3 and ZS97RS3, leaving 6.0 Mb unaligned to all three references, so most of this sequence reflects subspecies divergence and only a small part is specific to ZN65. Projection onto the 3,000 Rice Genomes panel placed ZN65 within *indica*: all 50 nearest accessions were *indica* (Additional file 2: Table S11).

**Figure 2.**
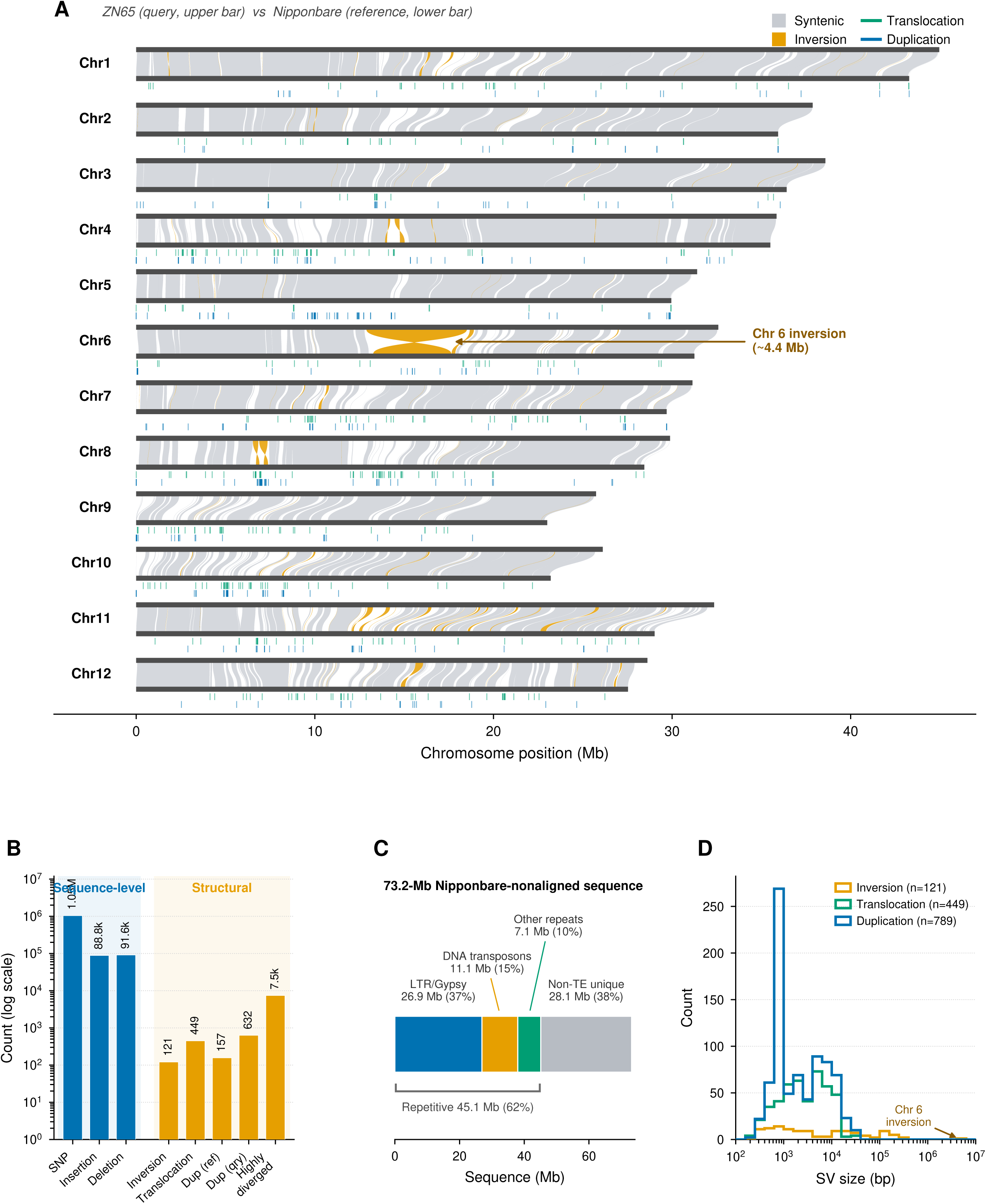
Genome-wide structural divergence of ZN65 from the *japonica* reference Nipponbare. (A) Whole-genome synteny (minimap2 asm5 + SyRI, visualized with plotsr): grey, syntenic; orange, inversions (note the large chromosome-6 inversion); green, translocations; blue, duplications. (B) Variant spectrum (log scale) split into sequence-level (SNP, insertion, deletion) and structural (inversion, translocation, duplication, highly diverged) classes; the 449 translocations shown are the SyRI aggregate of translocations (198) and inverted translocations (251). (C) Transposable-element families versus non-TE unique sequence within the 73.2 Mb of Nipponbare-nonaligned sequence (RepeatMasker). (D) Size distribution of structural variants. Robustness to minimap2 stringency and an orthogonal Assemblytics call are reported in Methods and Additional file 2: Table S3.

### Flavonoid pathway inventory and the pericarp pigmentation loci

KEGG orthology assigned 100 ZN65 genes to fifteen enzymatic steps of the flavonoid and anthocyanin pathway, from phenylalanine ammonia-lyase to the anthocyanidin 3-*O*-glucosyltransferase. Extension to the orthogroups of these genes gave 145 genes in ZN65 and 136 loci in Nipponbare (Figure 3A; Additional file 2: Table S4). Thus, we found no evidence of a pathway-wide expansion in ZN65. When the proteins of the 136 Nipponbare loci were aligned to the ZN65 genome, 114 matched an annotated ZN65 pathway gene and 12 matched another annotated gene. Ten had no annotated ZN65 gene at the aligned position: five with an intact reading frame, four of which encode proteins shorter than 121 residues, and five with a frameshift or incomplete coverage. One ZN65 gene model (ZN6511G1763) spanned 289,585 bp and covered seven Nipponbare loci (six adjacent loci on chromosome 11 and a chromosome-7 paralogue aligned to the same position). Therefore, the annotation contains a small number of missed and merged models in this pathway, and the gene counts are approximate. This result directs attention to the regulatory loci, although allelic or expression differences among the biosynthetic genes remain possible.

**Figure 3.**
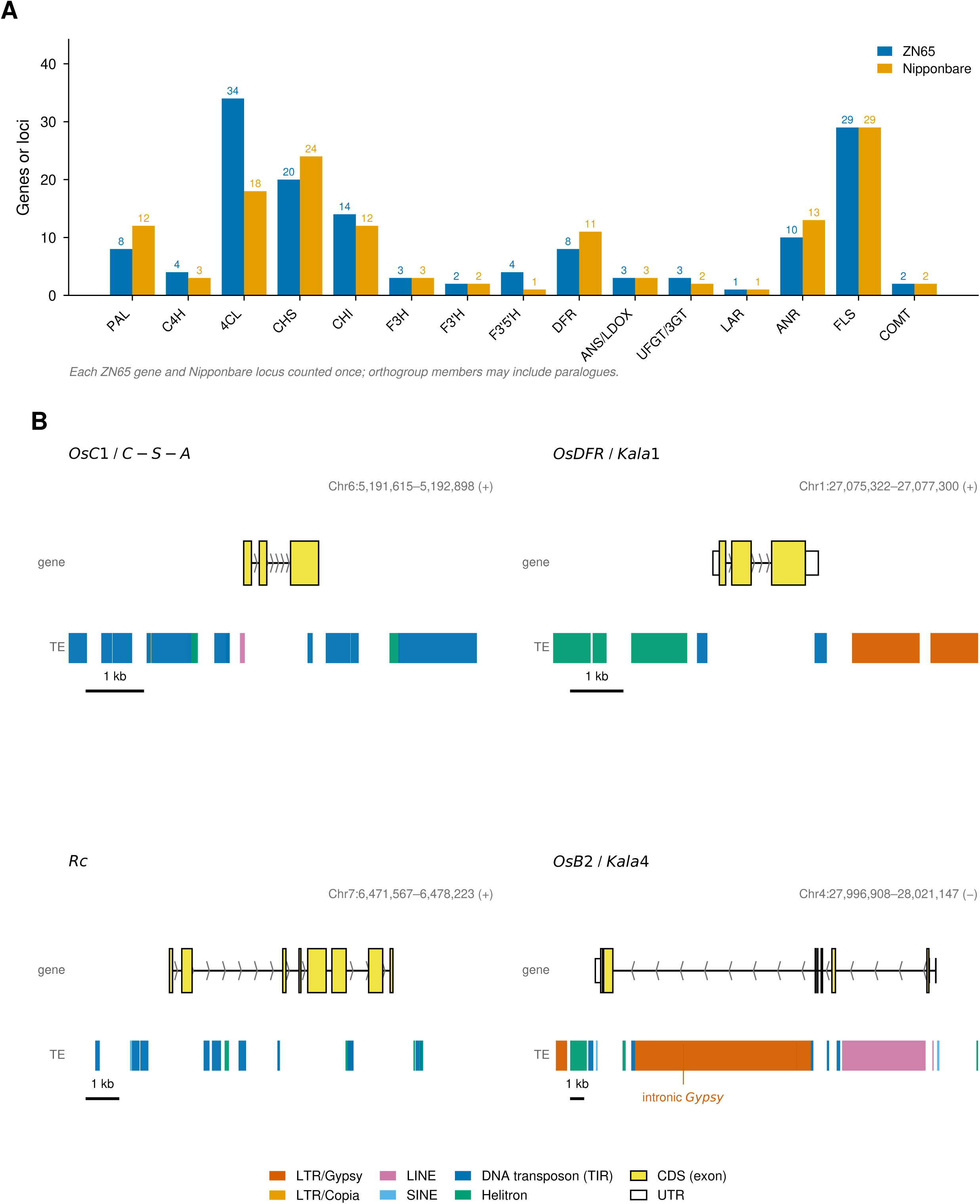
Flavonoid-pathway gene inventory and four pigmentation-related loci in ZN65. (A) Number of genes (ZN65, blue) and loci (Nipponbare, orange) assigned to each of fifteen enzymatic steps of the flavonoid/anthocyanin pathway by KEGG orthology and orthogroup membership, each gene counted once (totals 145 and 136). Anthocyanin end-products are not quantified. (B) Gene models (coding sequence, untranslated regions and introns) and RepeatMasker transposable-element annotation (coloured by class) of four pigmentation-related loci in ZN65, with coordinates above each panel: *OsC1* (chromosome 6; hull/apiculus *C*–*S*–*A* system), *OsDFR*/*Kala1* (chromosome 1), *Rc* (chromosome 7) and *OsB2*/*Kala4* (chromosome 4). *OsB2*/*Kala4* carries a ∼12.5-kb intronic *Gypsy*. Its upstream three-copy block is shown in Figure 4 and the *OsKala3* promoter repeat in Figure 6A. A 1-kb scale bar is shown per panel.

At the pericarp-pigmentation loci (Additional file 2: Table S7), ZN65 encodes apparently intact proteins at *OsKala3* (R2R3-MYB), *OsB2*/*Kala4* (a full-length 451-residue bHLH; the Nipponbare reference model is truncated at 180 residues) and *OsDFR*/*Kala1* (99.6% identity). Most diagnostically, ZN65 carries a full-length *Rc* (678 residues) with an intact bHLH domain, whereas white-pericarp Nipponbare carries the recessive *rc* allele, in which a 14-bp deletion in exon 6 truncates the C-terminal bHLH domain [29]. The WD40 partner *OsTTG1* (ZN652G3195) is full-length and 99.7% identical to its Nipponbare orthologue [34], so all three MBW components are present.

Structurally, *Kala1*/*OsDFR* (96.6% identity over 93.3% aligned length, with an ∼0.87-kb indel) and *Rc* (aligning over only ∼81% of the locus) carry transposon-associated variation, and *Kala4*/*OsB2* carries intronic retrotransposons and the upstream rearrangement described below (Figure 3B). *OsC1* (ZN656G0716) on chromosome 6, an R2R3-MYB that acts in the leaf sheath, apiculus and hull and is distinct from *OsKala3* [3], is colinear with Nipponbare at 99.8% identity. *OsKala3* (ZN653G2337) on chromosome 3 carries two directly abutting promoter RUs of 331 and 335 bp (94.3% identical) that span 701 to 36 bp upstream of the ATG, whereas the orthologous Nipponbare promoter carries one (Figure 6A). ZN65 thus carries the two-unit configuration reported for black-pericarp cultivars [3].

Two ancillary results are reported in Additional file 1. The chromosome-2 candidate originally nominated for this study, *Os02g0139500*, is *OsOSC1*, a 2,3-oxidosqualene cyclase of the triterpene/phytosterol pathway, and the chromosome-2 MYB ZN652G3275 clusters with lignin/secondary-wall MYBs and not with anthocyanin MYBs. Neither was considered further (Additional file 2: Table S9; Additional file 1: Figures S1 and S2).

### Q1: *Kala4*/*OsB2* carries a non-tandem three-copy upstream block supported by independent evidence

The *Kala4*/*OsB2* locus (ZN654G2687; chromosome 4: 27,996,908–28,021,147, minus strand; ∼24 kb) carries a complete ∼12.5-kb intronic LTR/*Gypsy* element, a 5,934-bp intronic long interspersed nuclear element 1 (LINE-1; LINE1-11_OS) and a promoter-proximal cluster of short interspersed nuclear element (SINE), Helitron, hAT and PIF-Harbinger elements (Figure 4B; Additional file 2: Table S6). The two LTRs of the *Gypsy* element differ by five substitutions over 994 ungapped sites (K2P distance 0.00505), giving an insertion age of ∼0.2 Myr (95% bootstrap CI 0.04–0.39 Myr). Thus, the insertion predates rice domestication. Its effect on *OsB2* expression remains untested.

**Figure 4.**
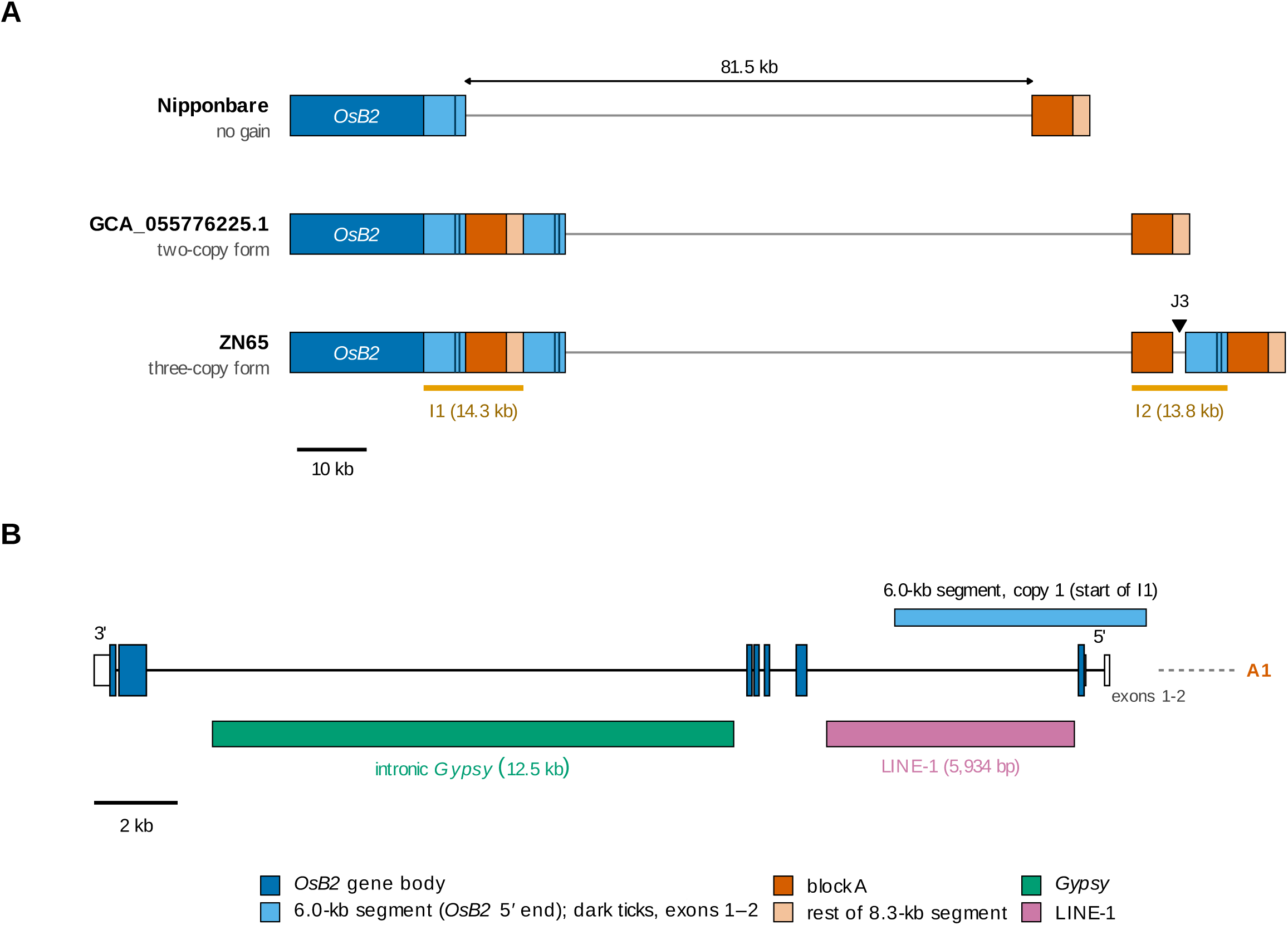
Three-copy upstream block and intronic retrotransposons at *Kala4*/*OsB2*in ZN65. (A) A ∼155-kb overview (chromosome 4, minus strand). Upstream of *OsB2*, a ∼5.87-kb block (A) occurs in three copies (A1–A3). The two extra copies lie within composite gains I1 and I2, which flank a collinear block (∼87.5 kb). The same block is a single homologous hit at the orthologous locus in the 33 other assemblies of the initial screen, including Nipponbare, *indica* (MH63, ZS97), *aus* (N22), wild *O. rufipogon* and the assembly registered as Cempo-Ireng (Additional file 2: Table S10). Three ultra-long Nanopore reads span the entire three-copy region (A1 to A3, 115,450 bp) as single molecules. The intronic LINE-1 element (LINE1-11_OS) spans the I1 junction and is marked there. (B) *OsB2* gene model with exon–intron structure (from the annotation): a complete *Gypsy* LTR retrotransposon (∼12.5 kb) sits within the largest intron and the 5,934-bp LINE-1 (LINE1-11_OS) within the gene. The *Gypsy* LTRs date the insertion to ∼0.2 Myr (K2P over 994 ungapped sites; 95% CI 0.04–0.39; µ = 1.3 × 10^−8^). The LTR identity shown in the panel (96.4%) is a whole-alignment value that includes indels.

Immediately upstream of *OsB2*, a ∼5.87-kb segment (block A) is present in three non-tandem copies (A1–A3). The two additional copies, A1 and A2, lie within composite segmental gains relative to Nipponbare (I1, ZN65 chr4:28,016,101–28,030,442; I2, 28,117,954–28,131,710), and A3 lies immediately distal to I2. The gains flank an ∼87.5-kb block that is collinear with Nipponbare (Figure 4A). It begins with the second copy of the 6.0-kb segment described below and continues ∼81.5 kb upstream. I1 consists of a copy of a 6.0-kb segment at the 5′ end of *OsB2* (Nipponbare chr04:27,934,782–27,940,807) followed by block A and 2.4 kb of its downstream sequence. The Nipponbare counterpart of this 8.3-kb part (chr04:28,022,289–28,030,604) lies 81.5 kb upstream of the gene segment. I2 contains block A, 987 bp of the same downstream sequence, an 871-bp interval and a third copy of the 6.0-kb segment. In ZN65 the 6.0-kb segment comprises the sequence between the proximal junction and the annotated transcript, the first two annotated exons (16 and 175 bp), the 451-bp intron between them and the first 4,402 bp of the following intron. Its Nipponbare counterpart contains exon 1 of transcript Os04t0557500-01 with the start codon. ZN65 carries three identical copies of this segment (6,027 bp): one contiguous with the rest of the gene (chr4:28,016,104–28,022,130), one distal to A1 (28,030,445–28,036,471) and one within I2 (28,125,684–28,131,710). Each differed from the Nipponbare segment at one of 6,026 aligned positions. The two gene-proximal copies flank an 8,314-bp segment (28,022,131–28,030,444) that contains A1, and an identical copy of that segment lies immediately distal to I2 and contains A3. Copies A1 and A3 were identical over block A and 2 kb on each side (9,870 bp), and A2 differed from them by one mismatch and one indel within the 5,870-bp block (the sequence shared by A2 extends 987 bp beyond the block). All three copies lie in the same orientation.

Several lines of evidence show that this structure is not an assembly artefact. HiFi coverage was continuous across the locus (minimum depth ≥11×); 6, 16 and 7 HiFi reads fully spanned I1, I2 and the intronic *Gypsy*, respectively; and three ultra-long Nanopore reads (MAPQ 60) spanned the entire three-copy region (A1 to A3, 115,450 bp) as single molecules (Additional file 2: Table S1). An assembly of the HiFi reads alone, which were held out of contig construction, placed the 250-kb region on one contig as a single alignment block, with three block A copies separated by 13,755 and 95,831 bp (13,756 and 95,824 bp in the final assembly). Inspector reported no structural error within chr4:27.9–28.2 Mb. When the ZN65 short reads were mapped to Nipponbare, the calibrated block A copy estimate was 2.70 and 23 reads were soft-clipped at the predicted junction base. No heterozygous SNP was called within chr4:27.99–28.16 Mb. Hi-C contacts between the copies fell within the background range (13th and 14th percentiles) and were uninformative within the copies, where reads cannot be placed uniquely.

To ask whether this configuration is shared among assembled genomes, we screened 34 distinct assemblies spanning the major cultivated groups, the assembly registered as Cempo-Ireng, wild *O. rufipogon* and wild and African outgroups (Additional file 2: Table S10). ZN65 gave three qualifying hits to block A, and each of the 33 other assemblies gave one. Of those 33 hits, 23 met both the 95% identity and the 95% coverage threshold (identity 96.31–99.98%, full query coverage). The other 10, mostly wild and African outgroups, were coverage- or divergence-limited. Therefore, block A is not novel sequence. What distinguishes ZN65 from these 33 assemblies is two additional, non-tandem copies. Because the single-hit state occurs in several cultivated groups and in the wild progenitor, the extra copies are most parsimoniously a derived gain. One junction coincides with the LINE-1 element (LINE1-11_OS), but the junctions lack the paired terminal repeats and target-site duplications of a complete transposon insertion, so we describe the structure as a LINE-associated complex segmental duplication.

The ZN65 configuration and the canonical black-pericarp *Kala4* allele [2] are compared in Table 1. The two descriptions agree in the distance between the gene and the source of the inserted segment (about 83 kb, and 81.5 kb from the end of the 6.0-kb segment in Nipponbare) and in the position of the insertion (954 bp and 984 bp upstream of the gene). They also have the same kinds of elements in the same order. The canonical allele was described with a duplication that includes exon 1, intron 1, exon 2 and part of intron 2 of *OsB2* (4,679 bp in the figure legend; the text describes it as extending from 4.9 kb upstream of exon 1 to within intron 2) and with an 11.02-kb insertion of a segment from about 83 kb upstream. In ZN65 the duplicated unit is the 6.0-kb segment, which includes the first two annotated exons, and the two gene-proximal copies are separated by the 8.3-kb segment, which consists of block A and 2.4 kb of its downstream sequence. The lengths differ between the two descriptions (4,679 and 6,027 bp for the duplicated unit, and 11.02 and 8.3 kb for the inserted segment). Oikawa et al. [2] classified black-rice promoters by the length of the inserted fragment, and their type II carries a deletion of about 2.7 kb at the 5′ junction. The 8.3-kb segment of ZN65 is close to the length expected for type II, but without sequence we do not assign ZN65 to a type. That study lists no sequence accession for the rearranged region, so the two structures cannot be compared base by base. Their correspondence is inferred from the description.

**Table 1.** The ZN65 upstream configuration compared with the canonical black-rice *Kala4* allele.

| **Feature** | **Canonical black-rice allele [2]** | **ZN65 (this study)** |
|---|---|---|
| Duplicated gene segment | 4679-bp proximal partial duplication spanning exon 1, intron 1, exon 2 and part of intron 2 | 6027-bp segment spanning the sequence from the proximal junction to the transcript, the first two annotated exons and 4402 bp of the following intron |
| Copies of the segment | Two | Three in ZN65; two in the three NCBI assemblies with the same junctions |
| Arrangement | Duplication accompanied by an 11.02-kb insertion of a segment whose donor lies about 83 kb away | The two gene-proximal copies flank an 8314-bp segment that contains block A and lies 81.5 kb from the gene segment in Nipponbare; the third copy lies within I2 with a further copy of block A |
| Position relative to the TSS | Insertion 954 bp upstream | Proximal junction 984 bp upstream of the annotated transcript 5′ end |
| Known variation | Three structural promoter types among 21 black-pericarp varieties, distinguished by deletions at the insertion ends | Two-copy and three-copy forms among 29 accessions with block A gain of 2913 surveyed |

TSS, transcription start site. For ZN65 the TSS is the annotated 5′ end of transcript ZN654G2687.1 (chr4:28021147, minus strand), not an experimentally mapped site, and the distance is the coordinate difference to the first base of the 8.3-kb segment (28022131) in the ZN65 assembly. The exons of ZN65 are those of transcript ZN654G2687.1, and the exon numbering of Oikawa et al. [2] may differ from it. Values for the canonical allele are from Oikawa et al. [2], who describe a representative canonical configuration.

A second screen covered the 210 *Oryza sativa* assemblies listed at NCBI and used stricter thresholds and junction probes (Methods). The control probe was present in 176 of the 210 assemblies. Of the 34 assemblies without it, 32 were contig-level. Among the 176, 143 carried one complete block A and no junction, 29 carried no complete block A hit and no junction, and one (Copsemar9, GCA_951799355.1) carried block A and the control probe on two chromosomes and no junction (Additional file 2: Table S14). Three assemblies carried the ZN65 junctions: Pulut-Hitam-2 (GCA_055776225.1, Malaysia), the assembly registered as Cempo-Abang (GCA_055776125.1, Indonesia) and Balatinaw (GCA_055776205.1, Philippines). Each carried two complete block A copies and the two I1 junctions, and lacked the A2-specific junction. The two copies were 95,834 bp apart in Pulut-Hitam-2 and 95,840 bp apart in the assembly registered as Cempo-Abang (95,824 bp between A1 and A2 in ZN65). The 135-kb ZN65 locus aligned to each of these two assemblies as one block in which the only difference of 500 bp or more was 13,755 bp present in ZN65 and absent from the assembly, within the I2 interval (I2 spans 13,757 bp). In Balatinaw the two copies were 89,862 bp apart, and the locus alignment showed the same 13,755-bp difference and two further differences between the copies (12,050 bp present only in Balatinaw and 17,928 bp present only in ZN65). The three assemblies therefore carry I1 without I2. In each of them the 6.0-kb segment was present in two copies that flanked the 8.3-kb segment, as in ZN65, and no third copy was found. The two copies were 99.98–100% identical to the ZN65 segment, and the 8.3-kb segment between them was 99.99% identical to that of ZN65 in all three assemblies (Additional file 2: Table S15).

Sedeek et al. [28] described Pulut Hitam 2, Balatinaw and Cempo Ireng as black and Cempo Abang and Zag as red. The assembly registered as Cempo-Ireng (GCA_055776245.1) carried a single block A hit (99.27% identity) and none of the junctions. Under the registered names, the two-copy form thus did not follow the reported colours. In addition, the assembly registered under the name of the red variety, Cempo-Abang, lacked the 14-bp *Rc* sequence required for a red pericarp [29], whereas the assembly registered under the name of the black variety, Cempo-Ireng, carried it. We therefore compared both assemblies with the reads of Sedeek et al. [28]. Of the first 20,000 reads deposited as Cempo Ireng, 19,256 aligned to the assembly registered as Cempo-Abang with a divergence of 0.47%, and 18,133 aligned to the assembly registered as Cempo-Ireng with a divergence of 4.19%. For the reads deposited as Cempo Abang, the corresponding values were 4.34% (18,405 reads) and 0.35% (19,255 reads). Thus, each read set matched the assembly registered under the other name. At the locus, the reads deposited as Cempo Ireng included 36, 40 and 37 reads spanning the three junction probes that were positive in the two-copy assemblies and covered block A at 1.97 times the flank depth, and none of the 58 reads spanning the *Rc* site carried the insertion. The reads deposited as Cempo Abang included no read that met the criteria for any junction probe or for the control probe and covered block A at 0.71 times the flank depth, and all 48 reads spanning the *Rc* site carried the 14-bp insertion. Therefore, the single-copy state of this genome rests on the matching assembly and on read depth. The read records are consistent with the reported colours at *Rc*, because the insertion required for a red pericarp [29] was found in the reads of the red variety. We followed the read records, although the public data leave open which record is in error. The reads of Pulut Hitam, Zag and Balatinaw matched the assemblies registered under their own names, with a divergence of 0.36–0.55% compared with 4.40–10.40% for the other four assemblies. With each assembly assigned to the variety whose reads matched it, the two-copy form was present in the three black varieties and absent from the two red varieties.

A third screen covered the 466 rice assemblies in the Genome Warehouse, which represent 264 materials (Methods; Additional file 2: Table S16). The control probe was present in all of them. In 175 materials the assemblies carried one complete block A and no junction. In 86 materials the best block A hit covered 94.4–94.5% of the query, just below the 95% threshold, and no junction was found; these hits reflect a block A haplotype with an indel of about 0.3 kb. Three materials carried the ZN65 junctions. MITAK (GWHESQM00000000), a japonica parent of an eight-parent population [35], and the Chinese *indica* accession Zaoshuxiangheimi (NH107; GWHBFLZ00000000 and the later version GWHDRDW00000000) carried the two-copy form. The ZN65 locus aligned to each of them as one block in which 13,755 bp (MITAK) or 13,925 bp (both NH107 versions) within the I2 interval were absent, and the segment identities of MITAK were the same as those of Pulut-Hitam-2. The third material, the Chinese *indica* accession Momi (NH202), was represented by two assemblies that differed at I2. In the chromosome-level version (GWHDRGW00000000), the ZN65 locus aligned without a break across I2 (64,490 of 64,600 bp between Chr4:28,078,916 and 28,143,514), and the 8.3-kb segment with block A was present in three copies. In the earlier contig-level version (GWHBFOZ00000000), the alignment broke within I2 (Chr4:28,125,077–28,129,836), although the contig had no gap in this region. The Nanopore reads of NH202 resolved the difference. All 25 reads that spanned both I2 anchors gave anchor distances of 20,769–22,090 bp (22,400 bp in ZN65), and none was close to the 8,643 bp expected without I2. All 17 reads that spanned both I1 anchors gave 19,770–20,751 bp (21,000 bp in ZN65). Distances on the reads were about 3% shorter than in ZN65 at both gains, as expected from deletion errors in Nanopore reads. NH202 therefore carries the three-copy form of ZN65, and the earlier assembly is incomplete at I2. The names Momi and Zaoshuxiangheimi refer to black rice in Chinese, but pericarp colour is absent from the sample records of the three materials.

The answer to Q1 is that ZN65 carries three non-tandem copies of block A, that this structure is supported by evidence independent of the assembly, that a two-copy form of the same structure is present in five public materials, and that the three-copy form is also present in the Chinese accession Momi, whose Nanopore reads span I2. In the gene-proximal part of the structure, two copies of the *OsB2* 5′ segment flank the 8.3-kb segment with block A. This part has the same kinds of elements in the same order as the description of the canonical black-rice allele, with different segment lengths.

### Q2: Block A gain occurs in 29 accessions in at least two copy-number forms and in 18 of 19 purple accessions

#### Frequency and copy-number forms

Of 3,011 accessions queried, 2,913 passed quality control (Additional file 2: Table S12). Read depth called 2,884 accessions (99.0%) at one copy, 12 at two copies and 17 at three or more (Figure 5A). Block A gain therefore occurred in 29 accessions (1.0%). The 17 accessions called at three or more copies were recorded from India (4), China (4), Indonesia (3), Myanmar (2), the Philippines (2), Laos (1) and Malaysia (1), and the 12 called at two copies from Indonesia (4), the Philippines (2), Thailand (2), Myanmar (2), China (1) and Malaysia (1). In 500-bp depth profiles, the accessions called at three or more copies had a median relative depth of 2.9–3.2 in the bins lying entirely within block A and the first 1 kb downstream of it, and 1.9–2.3 over the next 1.5 kb. The accessions called at two copies had a median of 1.7–2.2 in the bins within block A and the 2.4 kb downstream. Outside this interval the medians of both groups were 0.8–1.2 (Figure 5B). These profiles agree with the ZN65 structure, in which I1 contributes block A with 2.4 kb of downstream sequence and I2 contributes block A with 987 bp. Over the 6.0-kb segment at the 5′ end of *OsB2* (seven retained bins), the mean relative depth was 3.1–6.3 (median 3.7) in the accessions called at three or more copies, 1.6–3.2 (median 2.3) in those called at two copies and 0.4–1.7 (median 1.0) in the controls (Additional file 2: Table S15). Relative depth was also raised in the three usable bins next to the segment on the gene side (bin starts 27,933,000–27,934,000; medians 1.5–2.4 in the accessions called at three or more copies and 1.2–1.4 in those called at two), where ZN65 and the three assemblies carry one copy, so the duplicated interval of some carriers may extend beyond that of ZN65. Two accessions called at three or more copies (IRIS_313-10888 and IRIS_313-10950) differed from the rest: in 5-kb bins, relative depth was 1.5–2.5 in the 14 usable bins between the *OsB2* 5′ segment and block A (bin starts 27.940–28.015 Mb, upstream of *OsB2*) and 2.2–4.8 in the bins containing the *OsB2* 5′ segment and block A. This profile is compatible with an additional copy of the interval of about 96 kb from the *OsB2* 5′ segment to block A.

**Figure 5.**
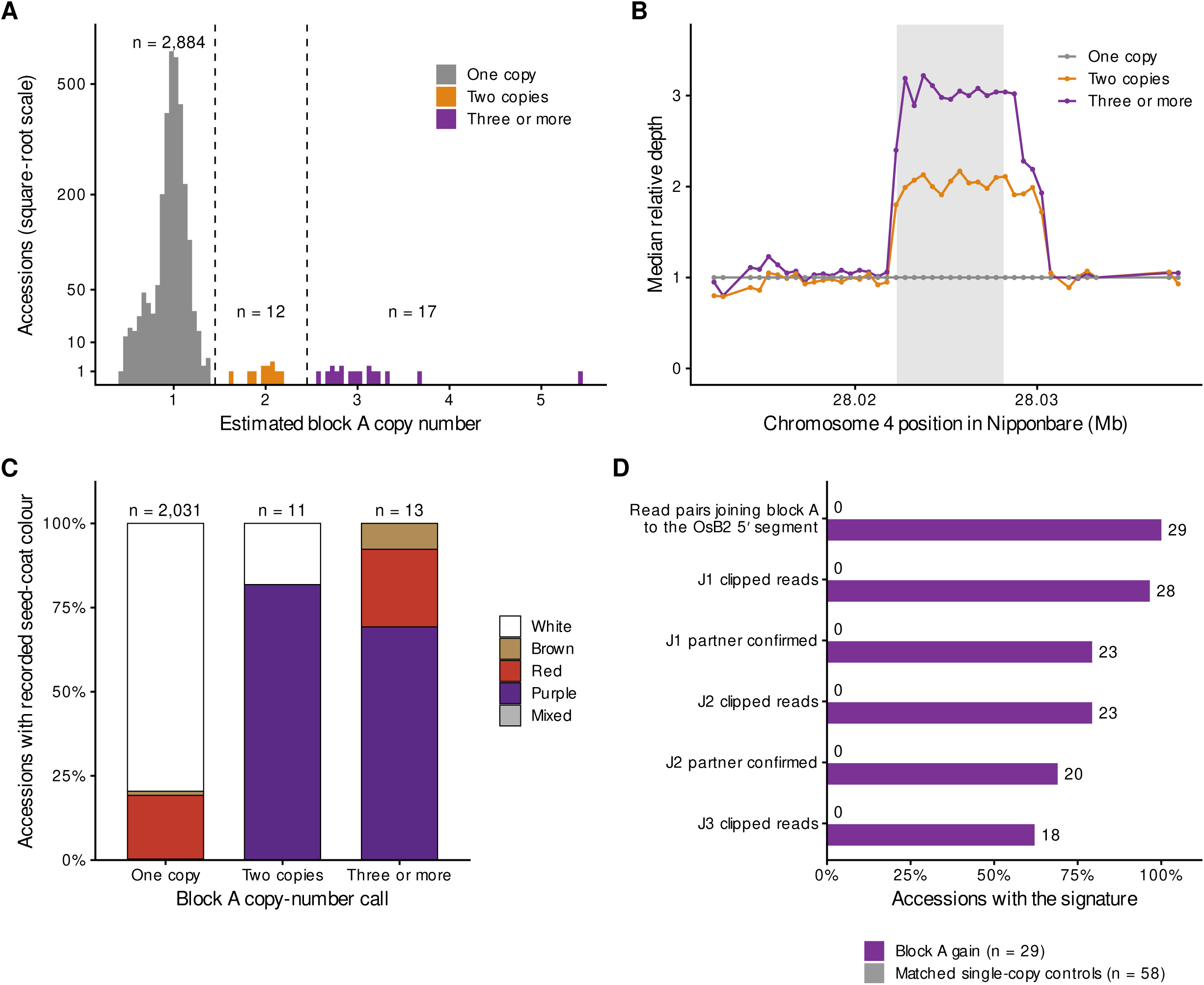
Block A copy number, junctions and seed-coat colour across 2,913 rice accessions. (A) Distribution of the estimated block A copy number, from read-depth ratios calibrated so that each batch median corresponds to one copy. Dashed lines, calling thresholds. (B) Median relative depth in 500-bp bins across chr04:28.012–28.038 Mb (IRGSP-1.0) for accessions called at three or more copies (n = 17), at two copies (n = 12) and for matched single-copy controls (n = 58). Bins in which the control median fell below 0.7 of the control density are omitted. Shading, block A. (C) Recorded seed-coat colour by copy-number call for the 2,055 accessions with a record. (D) The three junction signatures of the ZN65 structure in Nipponbare coordinates (J1, J2 and J3), with the number of carriers and controls in which soft-clipped reads and discordant read pairs were found.

#### Junctions

All 29 carriers had read pairs joining block A to the *OsB2* 5′ segment at 27.930–27.945 Mb (18–113 pairs), and none of the 58 matched controls had any (Additional file 2: Table S13). Reads soft-clipped at the J1 position were found in 28 carriers and at the J2 position in 23. The clipped segments mapped to the expected partner position in 23 and 20 carriers, respectively, and in 28 carriers for at least one of the two junctions. No control carried a clipped read at any of the three positions (Figure 5D). The A2-specific signature J3 was found in 16 of the 17 accessions called at three or more copies and in 2 of the 12 called at two copies (Fisher’s exact test, *P* = 3.3 × 10^−5^). Read-depth class and junction class therefore agree for 26 of the 29 carriers. Two accessions called at two copies had a heterozygous genotype fraction above 0.1 (IRIS_313-11817, which carried J3, and IRIS_313-11848), and a depth call of two may also arise from a heterozygous three-copy form. Without these two accessions, J3 was found in 16 of 17 and in 1 of 10 accessions (*P* = 2.0 × 10^−5^). Together with the public assemblies, these results indicate at least two forms, one that carries I1 without I2 and one that carries both. Read depth does not determine whether an individual accession is homozygous, and the structure of each carrier remains unverified with long reads.

#### Seed-coat colour

Seed-coat colour was recorded for 2,055 of the 2,913 accessions (Additional file 2: Table S12). Five of these records carried code 999 and were counted as not purple. Eighteen of the 24 carriers with a record were purple, three were red, one was brown and two were white. Both white carriers were called at two copies. Purple was recorded for 9 of the 13 accessions called at three or more copies and for 9 of the 11 called at two copies (Fisher’s exact test, *P* = 0.65), so the additional copy showed no relation to recorded colour. Of the 2,031 non-carriers with a record, one was purple (Figure 5C). Eighteen of the 19 purple accessions thus carried block A gain (two-sided Fisher’s exact test, *P* = 4.1 × 10^−38^). With code 999 treated as missing (n = 2,050) the test gave *P* = 4.3 × 10^−38^, and with the one purple record of code 088 also treated as missing (17 of 18 purple accessions with the gain) it gave *P* = 3.5 × 10^−36^. The association was also found within subpopulations (stratified exact test, *P* = 5.6 × 10^−33^; four informative strata), with ten principal components as covariates (Firth regression, *P* = 7.6 × 10^−20^; odds ratio for carriers about 1,350, 95% CI 287–6,377) and in a logistic mixed model with a genomic relationship matrix (score test, *P* = 2.5 × 10^−37^ without principal components and 4.1 × 10^−33^ with the first four). The mean pairwise relationship coefficient among carriers equalled that of random sets of non-carriers of the same subpopulation composition (0.023 in both). However, this comparison of means does not exclude close relatives among individual carriers. They do share the flanking region, as shown below, so these tests do not treat them as independent origins and the *P* values are exploratory.

#### Flanking genotypes

Within 200 kb of the locus (897 SNPs), the mean pairwise genotype distance among the 17 accessions called at three or more copies was 0.015, compared with a median of 0.205 (5th–95th percentile 0.181–0.227) for random sets of non-carriers with the same subpopulation composition. The median segment without opposite homozygous genotypes was 557 kb among these carriers and 12 kb in the random sets. ZN65 had a distance of 0.000 to the modal genotype of the carriers. The median distance to this genotype was 0.001 (range 0.000–0.046) for the 17 accessions called at three or more copies and 0.040 (0.004–0.173) for the 12 called at two copies (Additional file 2: Table S13). The 29 carriers belonged to tropical *japonica* (13), *indica* (6) and admixed (10) groups. Of 2,884 non-carriers, 276 lay within a distance of 0.02 of the carrier modal genotype, most of them tropical *japonica*, so a similar flanking genotype is common without the gain. Among accessions within this distance and with a colour record, 13 of 17 carriers and none of 169 non-carriers were purple. The answer to Q2 is that block A gain occurs in about 1% of the panel, that the carriers share the ZN65 junctions and fall into at least two copy-number forms, that the accessions called at three or more copies share a flanking genotype, and that the gain is carried by 18 of the 19 accessions recorded as purple.

### Q3: Co-occurrence of block A gain with an elevated *OsKala3* promoter-repeat dosage

Because the *OsKala3* RU is single-copy in Nipponbare, accessions carrying two RUs give about twofold read depth over it. The RU depth ratio was obtained for 2,899 of the 2,913 accessions, and 46 (1.6%) had a ratio of 1.5 or more, consistent with duplication of the RU (Figure 6B; Additional file 2: Table S12). However, RU copy number remains unverified at assembly level in these accessions. An elevated ratio was found in 21 of the 29 carriers of block A gain and in 25 of the 2,870 non-carriers (Fisher’s exact test, *P* = 3.0 × 10^−34^). The two variants therefore co-occur strongly, but eight carriers had no elevated RU ratio, and four of these eight were purple (Figure 6C). None of the 15 non-carriers with an elevated RU ratio and a colour record was purple. Among carriers with a colour record, 14 of 16 with an elevated RU ratio and 4 of 8 without were purple (*P* = 0.13). These relations held at thresholds from 1.3 to 1.5. At thresholds of 1.6 to 2.0 the number of purple carriers without an elevated ratio rose from 7 to 13 of 18, because the ratios of many carriers lie between 1.5 and 2.0. In a genome-wide linear mixed-model scan in which carrier status was included as one marker, carrier status ranked first, and the SNPs ranked next lay at chr03:16.88–17.17 Mb, the region of *OsKala3*. The answer to Q3 is that block A gain corresponds to purple seed coat more consistently than the RU depth ratio does, and that an independent contribution of *OsKala3* remains possible with these data.

**Figure 6.**
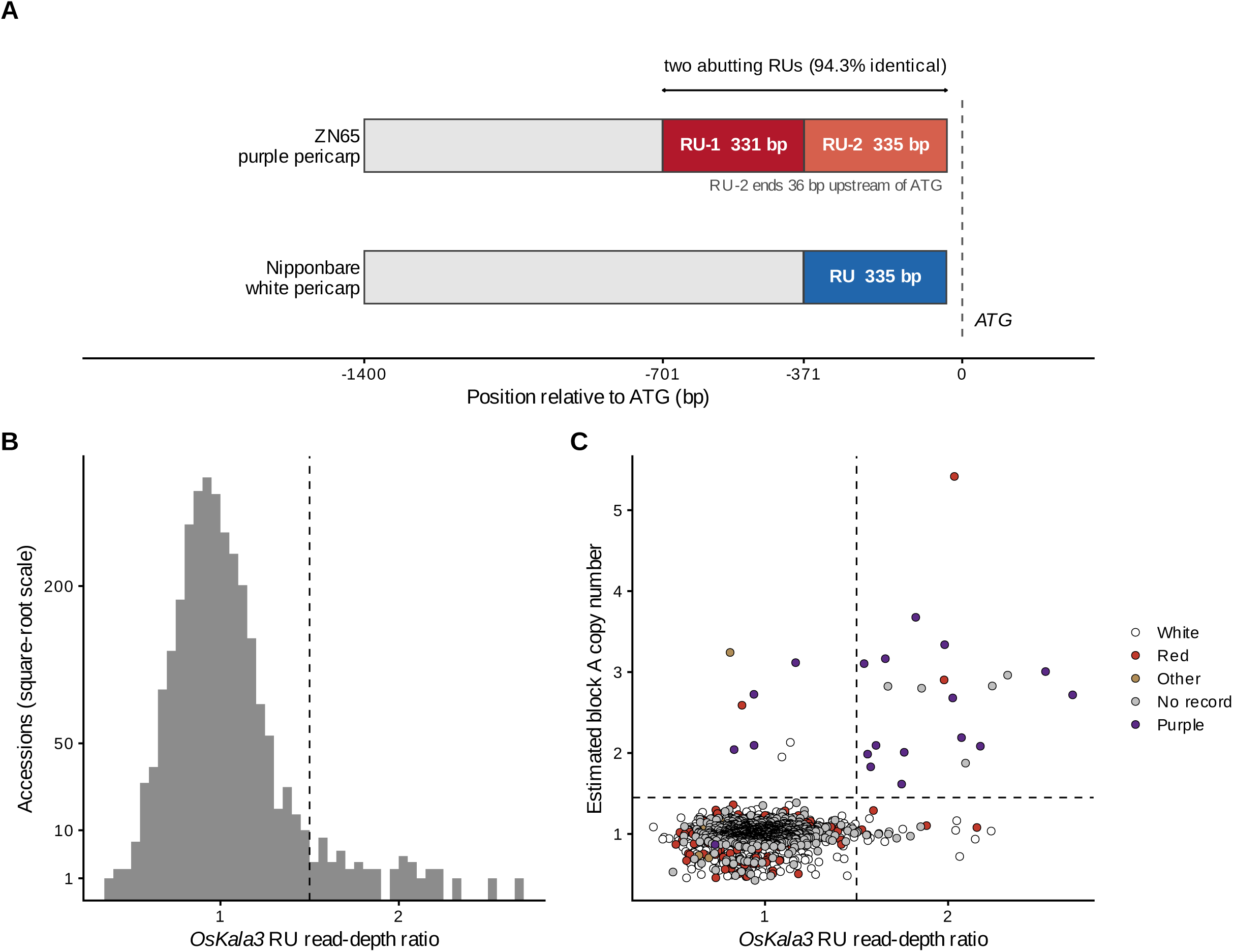
The *OsKala3* promoter repeat and its relation to block A gain. (A) Promoter architecture upstream of *OsKala3* (chromosome 3, minus strand). ZN65 carries two directly abutting repeat units, RU-1 (331 bp) and RU-2 (335 bp), 94.3% identical and spanning 701 to 36 bp upstream of the ATG (RU-2 ends 36 bp upstream of the ATG). The orthologous Nipponbare promoter carries a single 335-bp unit at the corresponding position. (B) Distribution of the *OsKala3* RU read-depth ratio across 2,899 accessions. Dashed line, the 1.5 threshold. (C) RU depth ratio against the block A copy estimate, coloured by recorded seed-coat colour. Dashed lines mark the RU threshold of 1.5 and the block A calling threshold of 1.45.

## Discussion

### Q1: a multicopy upstream structure resolved at sequence level

The canonical black-rice *Kala4* allele has been known from DNA gel blots and amplified fragments. The ZN65 assembly provides a gap-free sequence of a *Kala4*/*OsB2* upstream duplication that is supported by reads and by an independent assembly and, to our knowledge, the first sequence-level description of this rearrangement. The region carries three non-tandem copies of a 5,870-bp block. The structure was reproduced by an assembly of reads that were held out of contig construction, was spanned by single Nanopore molecules, and no structural error or heterozygous site was detected in it. A two-copy form with the same junctions is present in three of 210 public assemblies (registered as Pulut-Hitam-2, Cempo-Abang and Balatinaw), which are recorded as from Malaysia, Indonesia and the Philippines and were deposited by a single submitter. The Genome Warehouse added two materials with the two-copy form (MITAK and the Chinese accession Zaoshuxiangheimi) and one, the Chinese accession Momi, with the three-copy form. The chromosome-level assembly of Momi contains this form, which was previously undescribed. The canonical black-rice allele was described from DNA gel blots and amplified fragments as a partial duplication of the gene combined with an insertion of about 11 kb from a donor about 83 kb away [2]. The gene-proximal part of the ZN65 structure has the elements of that description in the same order: a duplicated segment of the *OsB2* 5′ end, an inserted segment whose other copy lies about 81.5 kb upstream in Nipponbare, and an insertion point about 1 kb upstream of the transcript. The same arrangement is present in the three NCBI assemblies at 99.98% identity or more, and read depth over the duplicated gene segment was raised in the carriers (medians of 2.3 and 3.7, compared with 1.0 in the controls). The lengths of the duplicated and inserted segments differ from the published values, and no sequence of the varieties studied by Oikawa et al. [2] is available. Correspondence with the canonical allele is therefore inferred from the description alone. ZN65 differs from Pulut-Hitam-2 and the assembly registered as Cempo-Abang by an additional 13.8-kb unit (their only difference of 500 bp or more) that carries a third copy of block A and of the 5′ gene segment, and Balatinaw differs further within the collinear block.

The registered names of the two Cempo assemblies and the read records of Sedeek et al. [28] were in a crossed correspondence, and the read records agreed with the reported colours at *Rc*. With each assembly assigned to the variety whose reads matched it, the two-copy form was found in the three black varieties of that study (Pulut Hitam 2, Balatinaw and Cempo Ireng) and was absent from the two red varieties (Cempo Abang and Zag). Which record carries the error remains unresolved with public data, and five varieties from one study are too few to establish an association with pericarp colour. The intronic *Gypsy* predates domestication, and whether it affects *OsB2* expression remains untested.

### Q2: two copy-number forms in purple-pericarp accessions

Block A gain was found in 1% of the panel, and all 29 carriers had read pairs joining block A to the *OsB2* 5′ segment, as in ZN65. Among accessions with recorded seed-coat colour, 18 of 19 purple accessions carried the gain. Block A gain with junction evidence matching ZN65 is therefore the predominant *Kala4*/*OsB2* upstream state of purple-pericarp accessions in this panel. However, the complete arrangement and zygosity of individual carriers remain undetermined. The gain is rare in the panel, as is purple pericarp. The accessions called at three or more copies shared a flanking genotype over several hundred kilobases and occurred in tropical *japonica*, *indica* and admixed groups. Black rice was proposed to have arisen in tropical *japonica* and to have spread to *indica* by introgression [2], and the distribution observed here is compatible with that history. The boundaries of introgressed segments were left unanalysed. A similar flanking genotype was found in 276 non-carriers, and none of the 169 with a colour record was purple. The order in which the flanking genotype and the gain arose cannot be determined from these data. The carriers fell into at least two forms that differed in the presence of the second copy, and in two accessions the depth profile was compatible with an additional copy of the whole 96-kb interval. Purple was recorded at similar proportions in both classes, so these data give no indication that I2 affects colour. The accessions called at two copies were recorded mostly from Southeast Asia, as were the three NCBI assemblies with this form, whereas eight of the 17 called at three or more copies were from India and China. Copy-number variation of this kind is expected at a locus with dispersed direct repeats, and its relation to the promoter types distinguished among black cultivars [2] remains unexamined.

### Q3: block A gain and the *OsKala3* promoter repeat

Duplication of the *OsKala3* promoter repeat has a published functional basis [3]. In this panel an elevated RU depth ratio co-occurred with block A gain, but it was not required for purple seed coat at the threshold used: four purple carriers lacked it, and none of the 15 accessions with an elevated ratio, without block A gain and with a colour record was purple. The RU interval is short and the ratio is imprecise near the threshold. The association signal on chromosome 3 near *OsKala3* indicates that purple accessions also share a genotype in that region. The present data cannot determine whether *OsKala3* contributes independently, because the two variants are strongly correlated and expression data are lacking.

### Limitations

Nine limitations bound these conclusions.

1. *Function.* The effect of block A on *OsB2* expression, and of RU dosage on *OsKala3* expression or promoter activity, remains untested. No pericarp transcriptome or anthocyanin measurement of ZN65 is reported.
2. *Phenotype.* Seed-coat colour was taken from germplasm records for 2,055 accessions without re-scoring. Six carriers were not recorded as purple, and anthocyanin content is unmeasured in all accessions.
3. *Non-independence.* The carriers share the flanking region, so the association tests do not represent independent events, and the effect of block A cannot be separated from that of linked variants. Only 19 accessions were purple, and the calibration of the score and Wald tests with so few cases remains unexamined.
4. *Copy number and arrangement.* Read depth gives relative copy-number estimates, not integer counts, and does not resolve zygosity. Junction signatures were detected with 83-bp reads, and clipped-read counts were taken without a mapping-quality filter. The structure of individual carriers, including the two accessions with elevated depth over the 96-kb interval, remains unverified with long reads or amplification across the junctions.
5. *Relation to the canonical allele.* No sequence of the allele described by Oikawa et al. [2] is available. The correspondence stated here rests on the published description and on three NCBI assemblies that were assigned by their reads to varieties described as black by Sedeek et al. [28], and the differences in segment length between that description and ZN65 are unexplained. Whether and which of the three copies of the *OsB2* 5′ segment are transcribed in the pericarp remains untested.
6. *Flanking genotypes.* The SNP data are unphased and pruned, and genotype distances do not establish identity by descent.
7. *Assemblies.* Each assembly represents a single genome, and the absence of a junction from an assembly may reflect its completeness in this region. The control probe was absent from 34 of the 210 assemblies, and three of 176 evaluable assemblies is not an estimate of allele frequency. The three junction-positive assemblies were deposited by one submitter. The sample records of the junction-positive assemblies omit pericarp colour, and the colours cited here are those reported for varieties of the same names. For the two Cempo assemblies, the colour is that of the variety whose reads matched the assembly, and the reads of the other three varieties matched the assemblies registered under their names. In the Genome Warehouse, three of 264 materials carried the junctions; their sample records omit pericarp colour, and the black-rice meaning of the names Momi and Zaoshuxiangheimi remains unchecked against the plants. The two NH202 assemblies differed at I2, and its form was decided from the reads. Assemblies deposited only in archives other than NCBI and the Genome Warehouse remain unscreened.
8. *Annotation.* The pathway inventory showed missed and merged gene models, and the annotation remains unexamined for such errors genome-wide.
9. *Assembly validation.* The read-based checks detect errors within their resolution and do not prove their absence. The heterozygosity estimates from the direct count and from the 21-mer spectrum differ, and amplification across the junctions of ZN65 is lacking.

### Next steps

Amplification across the junctions in ZN65 and in representative carriers, pericarp developmental expression of *OsB2*, *OsKala3* and *OsDFR* with anthocyanin measurement, and reporter assays of block A are required to convert this structural association into a causal account. Sequencing of the varieties in which the canonical allele was defined would settle its relation to the structure described here. The junction signatures can be used to track the structure in pigmented rice germplasm.

## Conclusions

A gap-free assembly of the purple-pericarp landrace Mojiang ZN65 gives the sequence of a *Kala4*/*OsB2* upstream duplication and shows that purple rice carries it in at least two copy-number forms. First, the region carries a three-copy, non-tandem arrangement of a 5,870-bp block that is supported by evidence independent of the assembly, a two-copy form with the same junctions is present in three of 210 NCBI assemblies and in two Genome Warehouse materials, and the three-copy form is present in the Chinese accession Momi. Second, block A gain was found in 29 of 2,913 accessions, all with read pairs joining block A to the *OsB2* 5′ segment, and in 18 of the 19 accessions recorded as purple. Third, an elevated *OsKala3* promoter-repeat depth ratio co-occurs with block A gain but is absent from four purple carriers. The gene-proximal part of the structure is shared with three NCBI assemblies and has the elements of the published canonical black-rice allele in the same order, with different segment lengths, and ZN65 carries an additional unit. The effect of the structure on *OsB2* expression remains to be established. The assembly and the junction signatures provide entry points for that work.

## Supplementary Information

Additional file 1 (.docx). Supplementary Methods and Supplementary Figures S1–S3. Additional methods for assembly and quality assessment, centromere and rDNA localization, sequence-periodicity analysis, comparative genomics, pathway inventory, transposon dating and subspecies placement; figure legends; and the legends of Tables S1–S16.

Additional file 2 (.xlsx). Supplementary Tables S1–S16. One worksheet per table; the legend of each table is given in Additional file 1.

## List of abbreviations

ATG: translation start codon
bHLH: basic helix-loop-helix
BUSCO: Benchmarking Universal Single-Copy Orthologues
CCS: circular consensus sequencing
CDS: coding sequence
CI: confidence interval
HiFi: high-fidelity long reads
K2P: Kimura two-parameter
KEGG: Kyoto Encyclopedia of Genes and Genomes
LAI: LTR Assembly Index
LINE: long interspersed nuclear element
LTR: long terminal repeat
MAPQ: mapping quality
MBW: MYB-bHLH-WD40
NOTAL: not aligned (SyRI category)
ONT: Oxford Nanopore Technologies
RU: repeat unit
SINE: short interspersed nuclear element
SNP: single-nucleotide polymorphism
T2T: telomere-to-telomere
TE: transposable element
TSS: transcription start site

## Declarations

### Ethics approval and consent to participate

Not applicable. This study did not involve human participants, human data, human tissue, or animals. The plant material used is a cultivated rice landrace described below; no endangered or protected species were involved, and all plant work complied with relevant institutional, national and international guidelines and legislation.

Mojiang purple rice (ZN65) is a traditional glutinous purple-pericarp rice landrace that originates from Mojiang Hani Autonomous County, Pu’er, Yunnan Province, China, where it has long been maintained by Hani custodian communities. The plants used in this study were grown at Pu’er University, Simao District, Pu’er, Yunnan, China (22°46′N, 101°58′E). Taxonomic identification was carried out by Fang Zhao (Pu’er University). A seed voucher of this accession is deposited in the seed collection of Pu’er University under accession number PR10010226. The genome libraries (PacBio HiFi, Oxford Nanopore ultra-long, Hi-C and DNBSEQ short-read) were prepared from genomic DNA of a single plant of this accession grown at the same site.

The material was obtained with the prior informed consent of the local custodian communities and is used exclusively for non-commercial academic research, in accordance with the Nagoya Protocol and China’s Regulations on the Administration of Access to and Benefit-Sharing of Biological Genetic Resources. The supporting documentation is retained by the authors and is available from the corresponding author on reasonable request.

### Consent for publication

Not applicable.

### Availability of data and materials

The raw sequencing reads, the telomere-to-telomere genome assembly and its annotation have been deposited at the China National Center for Bioinformation / National Genomics Data Center under BioProject PRJCA068452 (raw reads in the Genome Sequence Archive, accessions CRR3301760–CRR3301769; assembly and annotation in the Genome Warehouse), to be released publicly on publication. The 3,000 Rice Genomes Project alignments, genotypes and morpho-agronomic records are publicly available [26]; all other reference genomes are available under the accessions cited in Methods. Analysis scripts that regenerate the figures and tables are available on GitHub [36] and archived at Zenodo [37]. This manuscript is available as a bioRxiv preprint [38].

### Authors’ contributions

FeZ, Feng Zhao; FaZ, Fang Zhao. FeZ: Conceptualization, Methodology, Formal analysis, Investigation, Data curation, Visualization, Writing – original draft, Writing – review & editing, Supervision. JZ: Investigation, Formal analysis, Visualization, Writing – review & editing. FaZ: Investigation, Resources, Writing – review & editing. SB: Investigation, Resources, Writing – review & editing. YW: Investigation, Resources, Writing – review & editing. RZ: Investigation, Resources, Writing – review & editing. All authors read and approved the final manuscript.

## Acknowledgements

We thank the Hani custodian communities of Mojiang for the ZN65 landrace germplasm.

---

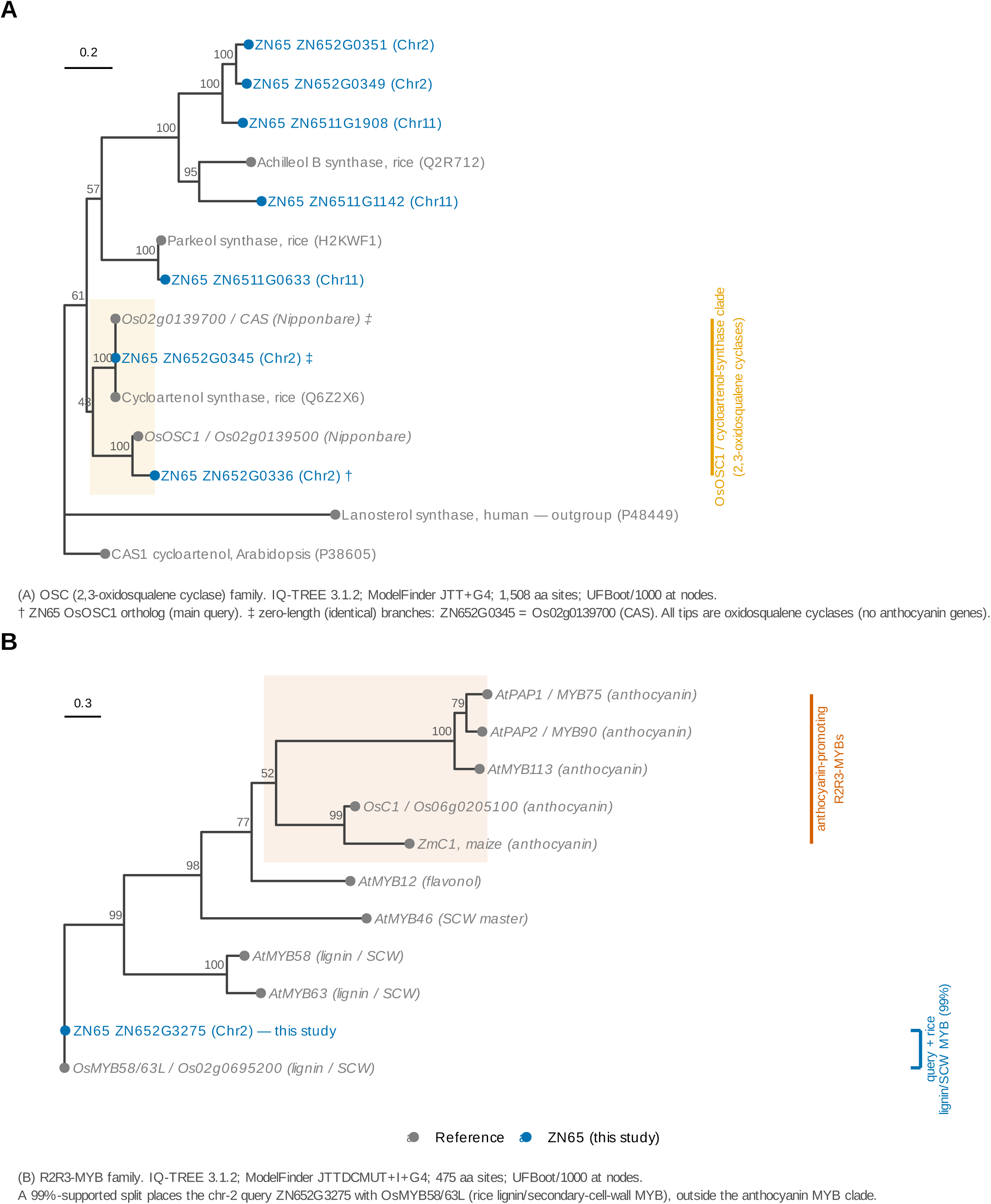
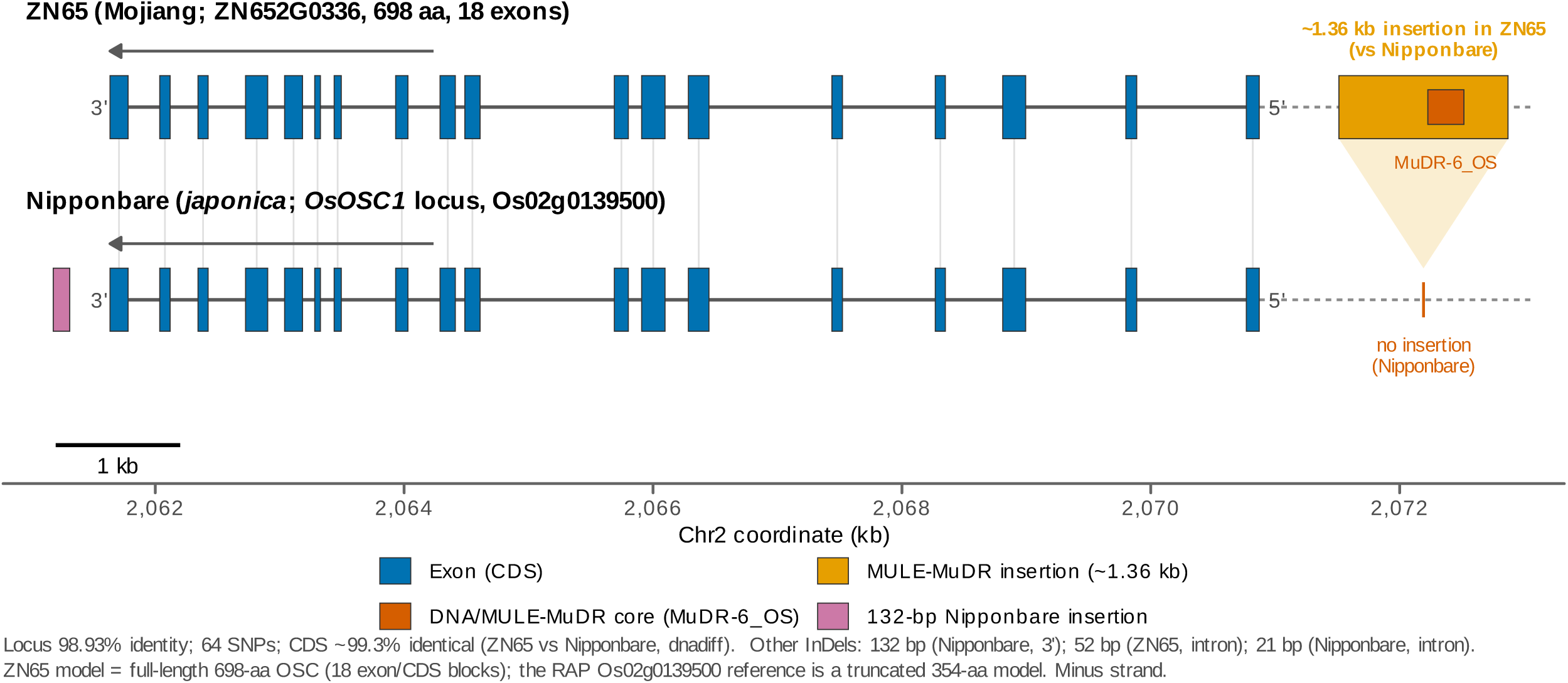
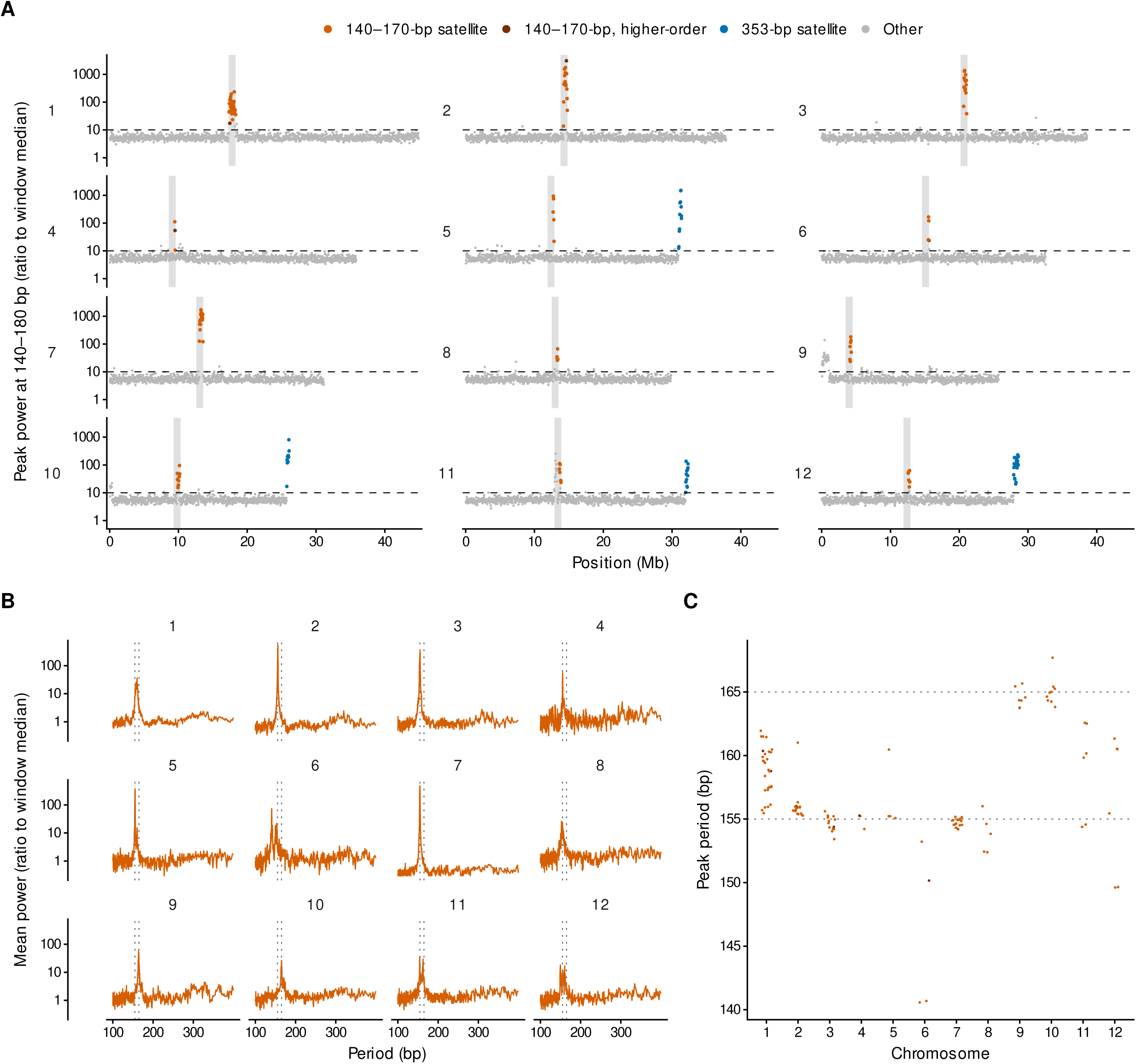

