## Supplementary Information (Supplementary Methods, Supplementary Figures S1-S3, table legends) for "Two copy-number forms of the *Kala4/OsB2* upstream duplication in purple rice revealed by a telomere-to-telomere genome of landrace ZN65"

### **Additional file 1**

This file contains Supplementary Methods, Supplementary Figures S1–S3 and the list of Supplementary Tables S1–S16 (provided as a separate spreadsheet, Additional file 2).

#### **Supplementary Methods**

**Assembly and quality assessment.** ZN65 was sequenced with PacBio HiFi (Revio; 20.6 Gb), Oxford Nanopore ultra-long (21.68 Gb), DNBSEQ (72.6 Gb) and Hi-C (39.3 Gb). Contigs (hifiasm v0.25.0, ONT mode) were polished (NextPolish; Winnowmap/bwa + Pilon), scaffolded with Juicer/3D-DNA, and gap-filled/telomere-completed with quarTeT v1.2.5. Quality was assessed with Merqury v1.3 (k = 19; QV 53.6, base error 4.4 × 10⁻⁶), the LTR Assembly Index (LTR_retriever; LAI 15.5) and BUSCO v5.8.0 (embryophyta_odb10; 99.6%, 1,607/1,614). Read-mapping rates were 99.96% (HiFi), 99.99% (ONT) and 99.31% (DNBSEQ).

**Centromere and rDNA localization.** Centromeres were identified as *CentO* satellite arrays: Tandem Repeats Finder monomers of 155–165 bp were clustered per chromosome, and the densest 1-Mb window per chromosome was taken as the centromere core (Table S5). The 45S (18S–5.8S–25S) and 5S ribosomal DNA arrays were localized from the BLASTn rRNA annotation by clustering adjacent rRNA features (gap ≤ 50 kb) into arrays and retaining arrays of ≥ 10 units spanning ≥ 20 kb (Table S5).

**Fourier analysis of sequence periodicity.** Each chromosome was scanned in windows of 65,536 bp with a step of 32,768 bp, and a final window was anchored at the chromosome end (12,053 windows). Each window was coded as four binary indicator sequences (A, C, G and T), each sequence was mean-centred, and the power spectrum was taken as the sum of the squared moduli of their discrete Fourier transforms (NumPy v1.26.4). Power was expressed as a ratio to the median of the window. As a reference, the bases of every tenth window were permuted (1,211 windows). The largest ratio in these windows was 8.6 at periods of 2–2,000 bp and 7.7 at 140–180 bp, and a ratio of 10 was used as a screening cut-off. For each window, the largest peak at periods of 140–180 bp was located and its period was refined by parabolic interpolation. Windows above the cut-off were classed in three steps. First, windows with a band period of 172.5–180 bp and a peak above the cut-off within 1.5% of twice this period were classed as a 353-bp satellite (59 windows). Second, windows in which the strongest peak at 185–2,000 bp exceeded the band peak were set aside as harmonics of a longer repeat (152 windows). These included every window above the cut-off that overlapped the 45S and 5S rDNA arrays of Table S5 (53 windows), and 14 further windows inside the centromere cores of Table S5 (band ratio at most 14.6). Third, the remaining windows with a band period of 140–170 bp were classed as a 140–170-bp satellite (123 windows). The *CentO* monomers of Table S5 fall in this range, but sequence identity was not tested for these windows. A window of this class was called higher-order when its strongest peak at 185–2,000 bp was above the cut-off and within 10% of two, three or four times the band period (6 windows). Windows of one class within 200 kb of each other were merged into runs, whose boundaries are therefore resolved to one step. The 140–170-bp windows formed twelve runs, one on each chromosome, and all runs overlapped the centromere cores of Table S5. Of these runs, nine lay inside the cores and three extended 11–50 kb beyond them (Figure S3). The distal boundaries of the 353-bp arrays were taken from the Tandem Repeats Finder records with a period of 350–359 bp in the terminal 1.5 Mb of the four chromosomes.

**Comparative genomics and structural variation.** Whole-genome alignment to Nipponbare (IRGSP-1.0) used minimap2 (asm5), SyRI and plotsr, with sensitivity checks at asm10/asm20 and an orthogonal nucmer/Assemblytics analysis (Table S3). Structural-variant counts are SyRI per-event-type annotations; the aggregate SyRI summary groups translocations with inverted translocations (449 = 198 + 251) and duplications with inverted duplications.

**Pathway inventory and orthology.** Genes of the flavonoid/anthocyanin pathway were assigned to fifteen enzymatic steps by KEGG orthology (100 ZN65 genes). Orthogroups between ZN65 and Nipponbare were inferred with OrthoFinder v2.5 (Diamond), and the set was extended to all ZN65 genes and Nipponbare loci in the orthogroups that contained a KEGG-assigned gene. Each orthogroup was assigned to one step, the step with the most KEGG-assigned ZN65 genes in it (ties to the step listed first in Table S4), and KEGG-assigned genes outside any orthogroup kept their KEGG step, so that each ZN65 gene and each Nipponbare locus was counted once (145 genes and 136 loci; Table S4). The proteins of the 136 Nipponbare loci were aligned to the ZN65 genome with miniprot, and each locus was classified by whether the aligned position overlapped an annotated ZN65 gene and whether the alignment was intact (query coverage of at least 0.9 with no frameshift or premature stop codon; Table S4).

**Transposon dating.** The intronic *Gypsy* element (RETRO2B) at *Kala4*/*OsB2* retains both LTRs (994 and 1,026 bp). The 5′ and 3′ LTRs were aligned (EMBOSS stretcher; 96.4% identity, 989/1,026), and over 994 ungapped sites differed by five point substitutions (indels excluded). The Kimura-2-parameter distance (0.00505) and the rice substitution rate µ = 1.3 × 10⁻⁸ site⁻¹ yr⁻¹ give *T* = *K*/(2µ) ≈ 0.2 Myr (95% CI 0.04–0.39 Myr by 1,000-replicate column bootstrap).

**Subspecies placement.** ZN65 was genotyped at 1,011,601 pruned 3K-RGP SNPs from its Nipponbare alignment, merged into the 3,024-accession panel (PLINK 1.9) and projected by PCA (PLINK 2.0); the 50 nearest accessions were all *indica* (Table S11).

#### **Supplementary Figures**

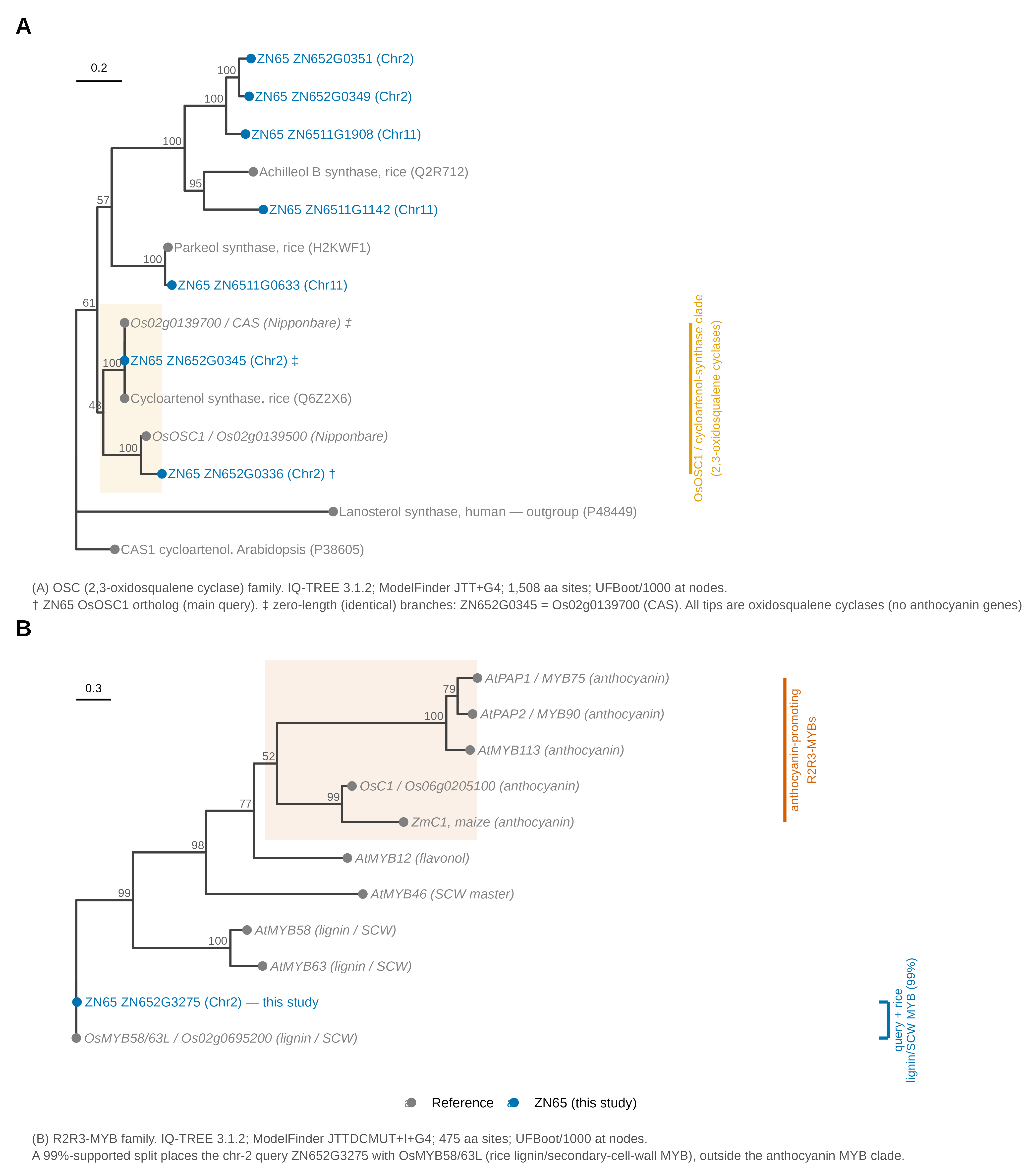

**Figure S1. The proposal gene *Os02g0139500* is *OsOSC1*, not an anthocyanin gene.** (**A**) Maximum-likelihood phylogeny (IQ-TREE 2, ModelFinder; ultrafast bootstrap) placing *Os02g0139500*/*OsOSC1* and its ZN65 orthologue (ZN652G0336) within the 2,3-oxidosqualene-cyclase (triterpene/sterol) family, distinct from anthocyanin biosynthetic genes. (**B**) Phylogeny placing the chromosome-2 MYB ZN652G3275 (= *Os02g0695200*) among lignin/secondary-cell-wall MYBs (AtMYB58/63 type), not the anthocyanin-regulatory MYB clade. Node values are UFBoot support; reference accessions are given at the tips.

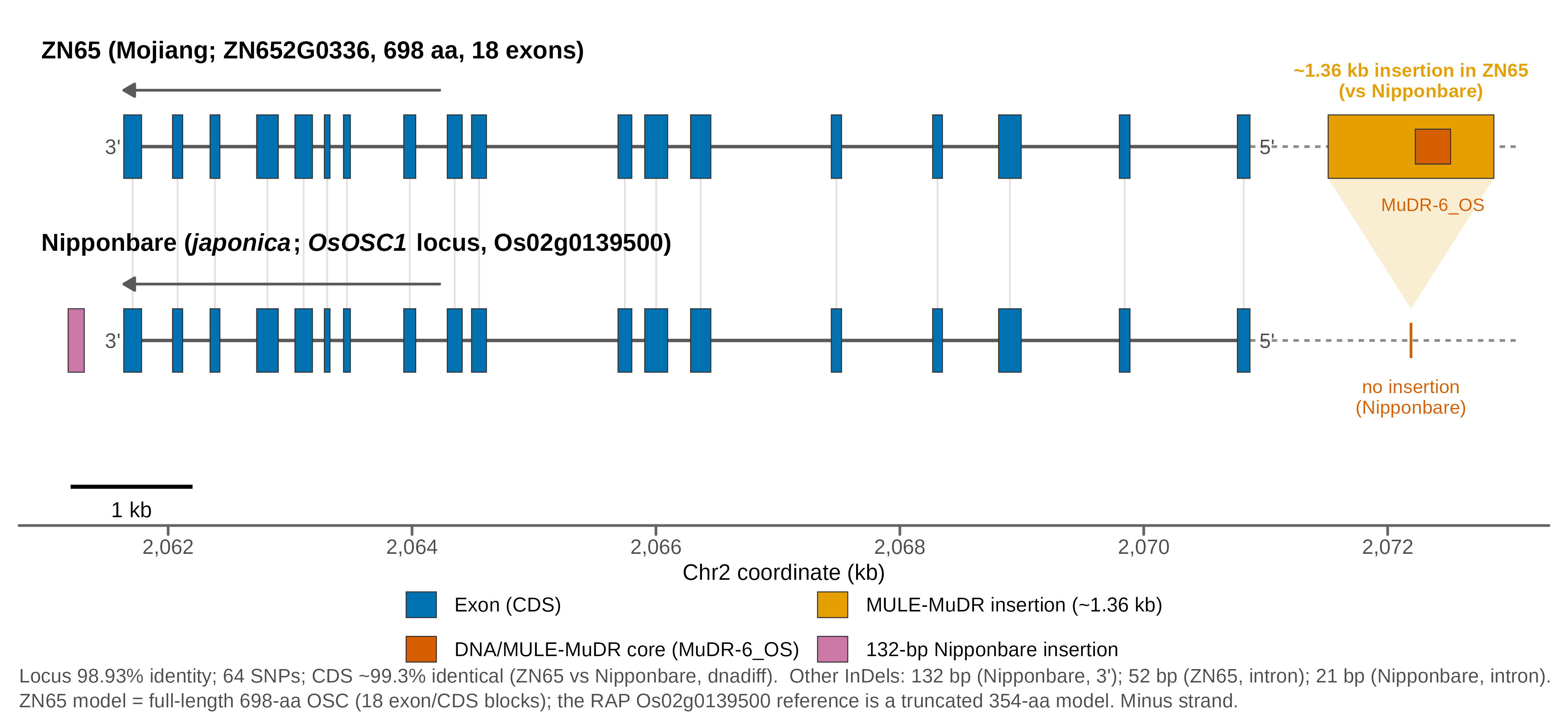

**Figure S2. Variant map of *OsOSC1* (*Os02g0139500*/ZN652G0336), ZN65 versus Nipponbare.** The 18-exon gene model with variants annotated: locus identity 98.93% (64 SNPs), CDS ~99.3% identical; the principal structural difference relative to Nipponbare is a ~1.36-kb MULE-MuDR insertion in the ZN65 5′ promoter (InDel sizes and TE families annotated).

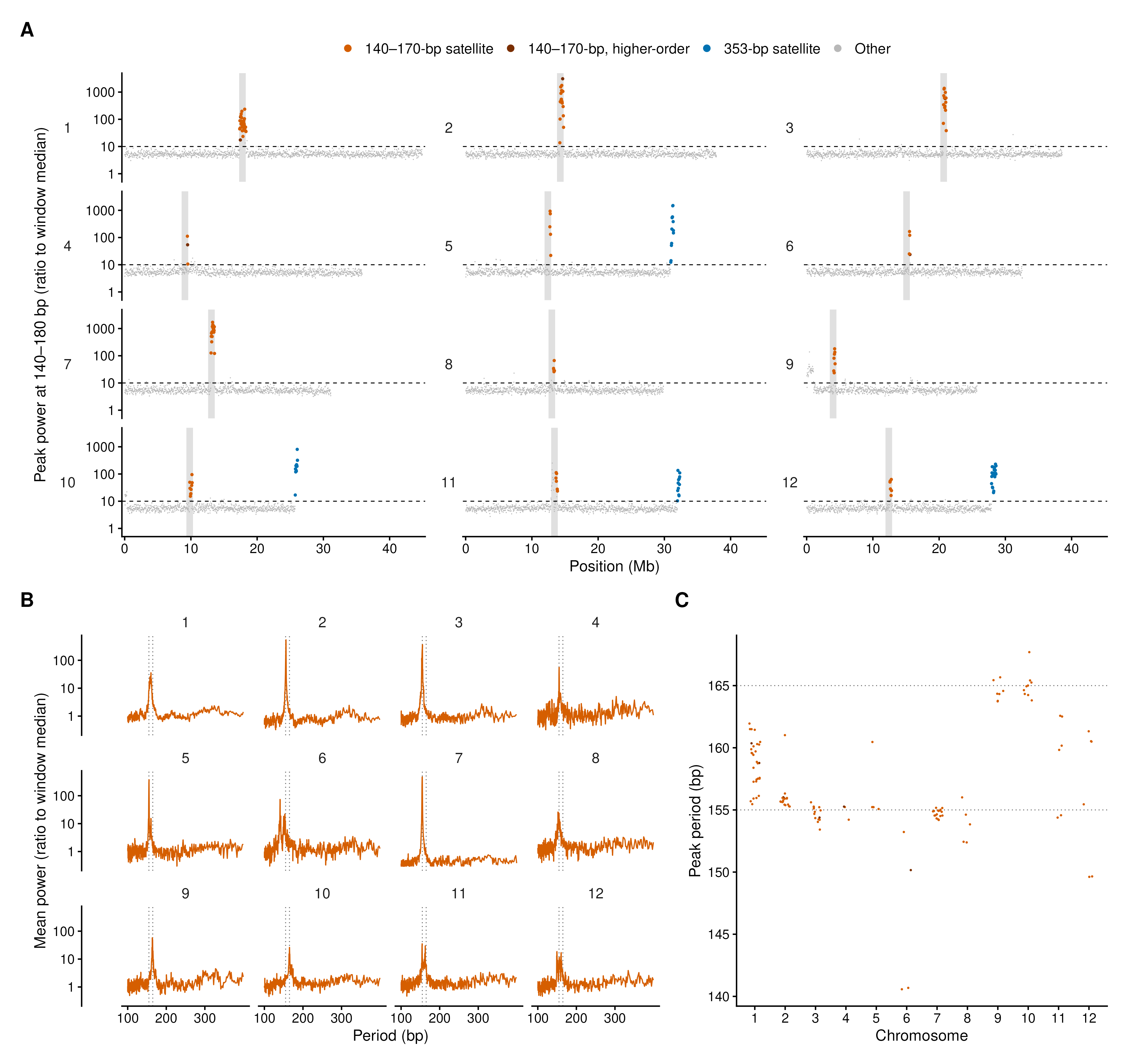

**Figure S3. Fourier analysis of satellite periodicity in the ZN65 assembly.** (A) Largest spectral peak at periods of 140–180 bp in windows of 65,536 bp along the twelve chromosomes, shown as a ratio to the window median. Windows are coloured by class, and higher-order windows are those whose strongest peak at 185–2,000 bp lay near a multiple of the band period. Windows in grey include those above the cut-off that were set aside as harmonics of longer repeats, such as the 45S rDNA at the start of chromosome 9 and the 5S rDNA on chromosome 11. Grey bands mark the centromere cores of Table S5 and the dashed line marks the cut-off of 10. (B) Mean power spectrum of the windows of the 140–170-bp run of each chromosome. Dotted lines mark 155 and 165 bp. (C) Peak period of each window of the twelve runs. The 353-bp satellite has a unit length similar to that of the 355-bp tandem repeat described by Ohtsubo et al. (1991, Jpn J Genet 66:241–254), and sequence identity was not tested.

#### **Supplementary Tables (Additional file 2)**

- **Table S1.** ZN65 T2T assembly statistics and quality metrics (per-chromosome length, telomeres, gaps, N50, GC, QV, LAI, BUSCO; long-read validation of the *OsB2* upstream segmental duplication and intronic retrotransposons).
- **Table S2.** Genome annotation summary (protein-coding genes, repeat classes, non-coding RNAs).
- **Table S3.** Genome-wide structural variation vs Nipponbare (SyRI; asm5/asm10/asm20 sensitivity).
- **Table S4.** Genes of the flavonoid/anthocyanin pathway in ZN65 and Nipponbare: genes and loci per enzymatic step with each gene and locus counted once, the 145 ZN65 genes, and the 136 Nipponbare loci with the result of aligning their proteins to the ZN65 genome.
- **Table S5.** Centromere (*CentO*) and 45S/5S rDNA array coordinates.
- **Table S6.** Transposable-element architecture of the *Kala4*/*OsB2* locus.
- **Table S7.** Functional-allele status at the four pericarp-pigmentation loci (ZN65 vs Nipponbare).
- **Table S8.** Gene nomenclature (symbol, *Kala* designation, RAP-DB locus, ZN65 ID).
- **Table S9.** Identification of the proposal gene *Os02g0139500* as *OsOSC1*.
- **Table S10.** Qualifying homologous hits to the *OsB2*-proximal ~5.87-kb block (A) in the initial screen of 34 distinct assemblies, one row per qualifying hit in each of the 36 genome files screened (three in ZN65 and one in each other file; two files are flagged as redundant inputs, leaving 34 distinct assemblies), with assembly accession, hit coordinates, query coverage, identity, merged-block count and QC status. The assembly registered as Cempo-Ireng (GCA_055776245.1) is listed with the variety assigned by its reads.
- **Table S11.** Subspecies placement of ZN65 (3,000 Rice Genomes projection).
- **Table S12.** Block A copy number in the 2,913 accessions of the 3,000 Rice Genomes panel that passed quality control (read counts in block A and in two single-copy control intervals, copy estimate and call, K9 group, seed-coat colour code and class, and *OsKala3* promoter repeat-unit depth ratio).
- **Table S13.** Junction evidence in the 29 accessions with block A gain and 58 matched single-copy controls (read pairs joining block A to the *OsB2* 5′ segment, clipped and through reads at junctions J1 to J3, heterozygous genotype fraction, flanking genotype distance and seed-coat colour).
- **Table S14.** Screen of 210 public *Oryza sativa* assemblies for block A and the ZN65 junction probes (hits per probe, class and positions of complete block A hits), with a note on the two assemblies whose registered names and read records were in a crossed correspondence.
- **Table S15.** Segments of the *Kala4*/*OsB2* upstream structure in ZN65 and four public assemblies (positions and identity of the 6.0-kb segment at the 5′ end of *OsB2*, the 8.3-kb segment, block A, I2 and the rest of *OsB2*), and relative read depth at the 5′ end of *OsB2* in the 29 accessions with block A gain and 58 matched controls.
- **Table S16.** Screen of 466 rice assemblies (264 materials) in the Genome Warehouse for block A and the ZN65 junction probes (assemblies, materials and classes), and Nanopore reads of NH202 that span both anchors of I1 and I2, with the anchor distances on each read.
